# Sucrose transport and metabolism control carbon partitioning between stem and grain in rice

**DOI:** 10.1101/2020.10.03.324889

**Authors:** Jyotirmaya Mathan, Anuradha Singh, Aashish Ranjan

## Abstract

The source-sink relationship is key to overall crop performance. Detailed understanding of the factors that determine source-sink dynamics is imperative for the balance of biomass and grain yield in crop plants. We investigated the differences in the source-sink relationship between a cultivated rice *Oryza sativa* cv. Nipponbare and a wild rice *Oryza australiensis* that show striking differences in biomass and grain yield. *Oryza australiensis*, accumulating higher biomass, not only showed higher photosynthesis per unit leaf area but also exported more sucrose from leaves than Nipponbare. However, grain features and sugar levels suggested limited sucrose mobilization to the grains in the wild rice due to vasculature and sucrose transporter functions. Low cell wall invertase activity and high sucrose synthase cleavage activity followed by higher expression of cellulose synthase genes in *Oryza australiensis* stem utilized photosynthates preferentially for the synthesis of structural carbohydrates, resulting in high biomass. In contrast, the source-sink relationship favored high grain yield in Nipponbare via accumulation of transitory starch in the stem, due to higher expression of starch biosynthetic genes, which is mobilized to panicles at the grain filling stage. Thus, vascular features, sucrose transport, and functions of sugar metabolic enzymes explained the differences in the source-sink relationship between Nipponbare and *Oryza australiensis*.

**Highlight:** Vascular features, sucrose transport, and sugar metabolic enzyme activity contribute to the differential source-sink relationship between the selected cultivated and wild rice that differ in biomass and grain yield.

## Introduction

Source-sink relationship is critical for determining the overall growth and performance of crop plants (White *et al*., 2016). Photosynthetic carbon assimilation in source leaves, sucrose transport through the vasculature, type and strength of sink tissues, as well as the metabolic status of the tissues are the key features that determine the source-sink dynamics of a plant. Photosynthetic leaves are the major source tissues of a plant, whereas types and strength of sink organs vary depending upon the growth stages and environmental conditions. For example, grains are the primary sink in the reproductive stage of crop plants, while roots, stem/internodes, and growing leaves function as the sink in the vegetative stage. Natural genetic variation for differences in the source-sink dynamics, resulting in changes in biomass and yield, has been evident (Yin *et al.*, 2009; Burnett *et al.*, 2016; White *et al.*, 2016; Fabre *et al.,* 2020). For example, many of the wild relatives of rice accumulate high biomass with poor grain filling, whereas cultivated varieties produce higher grain yield (Sanchez *et al.*, 2013).

Leaf morphological and anatomical features along with photosynthesis per unit leaf area largely determine the source strength of a plant (Mathan *et al.*, 2016). Functions of the key carbohydrate metabolic enzymes, such as ADP-glucose pyrophosphorylase (AGPase), sucrose phosphate synthase (SPS), sucrose phosphate phosphatase (SPP), and sucrose synthase (SUS), are known to be key determinants of the metabolic status of source leaves (Osorio *et al.*, 2014; Ruan, 2014). Similarly, the capacity to utilize photosynthates towards storage and maintenance largely determines the sink strength. Sink size, influenced by tiller number, spikelet number per panicle, and seed size; and sink activity, determined by functions of various metabolic enzymes such as cell wall invertase (cwINV) and sucrose synthase (SUS), dictate the photosynthates utilization at a sink organ (Smith *et al.*, 2018; Stein and Granot, 2019). Photosynthates are primarily partitioned from source leaves to different sink tissues in the form of sucrose via phloem with the aid of SWEET and SUT transporters (Braun, 2012; Chen *et al.*, 2012; Julius *et al.*, 2017). Mutations or overexpression of genes encoding sucrose transporters affect plant yield and biomass via modulation of source-sink dynamics (Scofield *et al.*, 2002; Chen *et al.*, 2012; Yang *et al.*, 2018). Besides the sucrose transporter functions, vascular features not only determine the rate of sucrose export from leaf tissues to phloem sap but also control mobilization of photosynthates to the panicles at the reproductive stage (Qi *et al.*, 2008; Fujita *et al.*, 2013; Sack and Scoffoni, 2013).

Photosynthates assimilated in leaves are either used to meet immediate cellular needs or mobilized into different sinks. Starch stored in leaves during the day is degraded during the night to be transported to different sink organs for growth (Smith and Stitt, 2007; Chen *et al.*, 2012). Mutants defective in sucrose transport, usually, accumulate starch in leaves at the end of the night, limiting plant growth (Chen *et al.*, 2012). The utilization and storage of sugars are regulated diurnally as well as developmentally. Stem serves as the main storage organ for carbohydrates in crop plants, including rice, during vegetative growth. Sucrose, after getting unloaded into the stem, could possibly be converted to different non-structural carbohydrates (NSCs), such as glucose, fructose, and starch. Starch is reported to be the major storage form in rice stem during the vegetative stage, which is mobilized to developing grains with the onset of the reproductive stage, contributing up to half of the final grain yield (Wang *et al.*, 2017). A portion of the photosynthates is also used as a structural component of the cell wall for the growth and development of new organs as well as elongation and thickening of existing organs. High biomass accumulation is usually associated with the accumulation of cellulose and other structural carbohydrates in crop plants, with cellulose alone contributing around 25-50% of plant biomass (Haigler *et al.*, 2001). Thus, the balance of structural and non-structural carbohydrates derived from the sucrose in the stem is critical for determining biomass and grain yield in cereal crops.

Sucrose, after reaching the stem, is hydrolysed by two classes of enzymes, invertases (INV) and sucrose synthase (SUS). INV cleaves sucrose into glucose and fructose, while SUS reversibly catalyses the formation of UDP-glucose and fructose. The UDP-glucose, generated by the cleavage activity of SUS, is used as a precursor for cellulose synthesis. Further, UDP-glucose can be converted to ADP-glucose for usage in starch synthesis, making SUS a key enzyme for biomass and grain yield (Smith *et al.*, 2012; Ruan, 2014; Stein and Granot, 2019). The functions of SUS are important for phloem loading and unloading as well as for determining sink strength (Smith *et al.*, 2012; Fan *et al.*, 2017; Yao *et al.*, 2020). Among the seven genes encoding SUS enzymes in rice, *OsSUS1* and *OsSUS3* contribute to seed starch biosynthesis (Cho *et al.*, 2011). Higher expression of *OsSUS3* also led to increased levels of structural carbohydrates, cellulose and hemicellulose, in rice (Fan *et al.*, 2017). *OsSUS1* is highly expressed in rice internodes, with a strong correlation with the expression pattern of genes encoding cellulose synthases, *OsCES4*, *OsCES7*, and *OsCES9* (Hirose *et al.*, 2008; Guevara *et al.*, 2014). OsCES4, OsCES7, and OsCES9 enzymes have been reported to function in secondary cell wall formation (Tanaka *et al.*, 2003; Wang *et al.*, 2016). Thus, functions of enzymes, such as INV, SUS, and CES, are important for determining source-sink dynamics for biomass and grain yield.

Despite the key importance of sucrose partitioning and metabolism in determining the source-sink relationship, this aspect is largely unexplored for optimization of grain yield and biomass in rice. Cultivated rice varieties and their wild relatives, with striking differences in yield and biomass traits, provide an excellent system to investigate the mechanistic basis of the source-sink relationship. Comparative characterization of source and sink features along with physiological and biochemical attributes for the selected wild and cultivated rice, which differ in grain yield and biomass traits, might provide key information related to bottlenecks for higher grain yield. In addition, such a study would also underline the factors contributing to high biomass in wild rice. The knowledge can be utilized for streamlining the sucrose transport and metabolism towards optimizing source-sink dynamics for higher yield and/or biomass. Here, we investigated the limitations related to sucrose transport and metabolism for grain yield in wild rice *O. australiensis*, resulting in higher biomass, compared to the cultivated variety *O. sativa* cv. Nipponbare. We report that a higher accumulation of structural carbohydrates, cellulose and hemicellulose, in *O. australiensis* is due to lower cell wall invertase activity and higher SUS cleavage activity together with higher expression of genes encoding cellulose synthases. Cultivated rice Nipponbare, in contrast, accumulated more starch in the stem due to higher expression of genes encoding starch biosynthesis enzymes. We also established the contribution of specific sucrose transporters and vascular features towards limited mobilization of photosynthates to *O. australiensis* panicles. Results from additional cultivated and wild rice accessions, *O. sativa* cv. IR 64 and *O. latifolia*, further corroborated our model.

## Materials and methods

### Phenotyping of the selected rice accessions for biomass and yield-related traits

Two cultivated varieties, *Oryza sativa* ssp. *indica* cv. IR 64 and *Oryza sativa* ssp. *japonica* cv. Nipponbare, and three wild rice species, *Oryza rufipogon* (IRGC 99562)*, Oryza latifolia* (IRGC 99596), and *Oryza australiensis* (IRGC 105272) were investigated for the study. Seeds were germinated on the germination paper in a petri dish for 5-7 days. Germinated seedlings were grown in half-strength Yoshida media for two weeks, and then transplanted in the experimental field of National Institute of Plant Genome Research, New Delhi (*latitude* 28°36 N; *longitude* 77°12 E; *altitude* 216 m). Plants were grown under natural growing conditions with average air temperature >30°C, 70-80% humidity, and more than 100 cm annual rainfall. All the phenotyping was performed after the transition to the reproductive stage. At least ten individual plants of each genotype were used for phenotyping. Plant architectural traits were quantified at the milk-stage of grain-filling. Seed traits and fresh and dry weight were quantified at the grain maturity stage. Plant height was measured as the height from the soil surface to the tip of the central panicle; total leaves per plant, tiller number per plant, and internode number per main stem were counted manually; and internode length was calculated using the distance between two consecutive nodes on the main stem. Leaf surface area was quantified for fully expanded top four leaves, including flag leaf, using a portable leaf area meter (LI-3000C, LI-COR Biosciences). Stem thickness and stem circumference were quantified from images of hand-cut sections of the main stem using Fiji-Image J software (Schindelin *et al.*, 2012). Fresh weight was quantified by directly weighing the entire plant mass, and dry weight was quantified by weighing the dried plant mass after drying the entire plant mass at 60°C for five days. The number of spikelets per panicle was counted manually. For grain weight analysis, 1000-grains were randomly selected and weighed using an electronic balance. For grain yield per plant, physiologically dried grains per plant were harvested and weighed. Raw data of biomass and yield-related traits for each genotype were normalized to the median, followed by log base 2 transformation for principal component analysis using JMP software from SAS (https://www.jmp.com/en_us/home.html).

### Quantification of leaf photosynthesis rate and related physiological traits

Leaf photosynthesis rate (*A*), stomatal conductance (*g_s_*), and intercellular CO_2_ concentration (*C_i_*) were recorded from flag leaves at booting, milk- and dough- stage of grain filling for *O. sativa* ssp. *japonica* cv. Nipponbare and *O. australiensis* using a portable photosynthesis system LI-6400XT (LI-COR Biosciences). *A*, *gs*, and *C_i_* were quantified from the flag leaf regions fully exposed to sunlight at the booting stage, as leaves were not fully expanded. The values were measured from the middle widest part of fully expanded flag leaves at the milk- and dough- stage of grain filling. Data were recorded from at least 15 independent plants of each genotype on a clear day between 9.00 h to 11.00 h at ambient CO_2_ level (*C_a_*; 400 μmol mol^−1^), photosynthetic photon flux density (PPFD, 1,500 μmol m^−2^ s^−1^), 300 μmol s^−1^ flow rate, and 70% relative humidity. During measurements, the leaf chamber air temperature was set at 30°C with maximum vapor pressure deficit in the range of 1.0-1.5 kPa.

### Quantification of glucose, fructose, and sucrose

The non-structural carbohydrates (NSCs: glucose, fructose, and sucrose) present in the flag leaf and stem were quantified at the milk-stage of grain-filling by Gas Chromatography-Mass Spectrometry (GC/MS). Flag leaves and stem tissues of both the genotypes were harvested from field-grown plants at 9:00-11:00 am. For the quantification of NSCs in flag leaves at the end of the day (EOD) and end of the night (EON), tissue samples were collected at 6.00 pm and 6.00 am, respectively. The tissues were sampled from at least four individual plants per genotype and immediately frozen into liquid nitrogen after harvesting. A 100 mg fresh weight (FW) of the tissues was extracted using 1.0 ml water: chloroform: methanol (1:1:2.5) spiked with 10 μl internal standard (1 mg ml^−1^ ribitol in water). The mixture was vortexed, centrifuged at 13,000 x g for 15 min., and 200 μl of polar phase was dried for 3 hours in a speed-vac concentrator. Next, dried samples were derivatized with 50 μl of 20 mg ml^−1^ methoxyamine in pyridine warmed for 37°C for 120 mins with shaking. Metabolites were further derivatized by adding 70 μl of N-methyl N-trimethylsilyl-trifluoroacetamide (MSTFA) at 37°C for 30 min. Once derivatization was completed, samples were transferred to GC compatible vials, and 1μl of samples was injected on GC-MS-QP2010 (Shimadzu) as described in Lisec *et al.* (2006). For metabolite identification, the mass spectra were matched with commercially available NIST spectral libraries, and the relative amount of the metabolites was calculated by the total ion current signal that was normalized to ribitol and tissue weight (Kim *et al*., 2012; Mikaia *et al*., 2014).

### Starch quantification

Starch was quantified from flag leaves, stems, different internodes, and mature seeds of the selected wild and cultivated accessions. The quantification of starch from flag leaves and stems was performed at the milk-stage of grain-filling at the time of phenotyping of photosynthesis. In addition, flag leaves samples were used for starch quantification at EOD and EON. Starch was also quantified from matured leaves at the vegetative and booting stage, as well as in different internodes of *O. sativa* ssp. *japonica* cv. Nipponbare and *O. australiensis* at the milk-stage of grain-filling. Starch was quantified using Mega-Calc Total starch determination kit (K-TSTA; Megazyme) according to the manufacturer’s protocol. Four independent biological replicates were analysed for each tissue type per genotype.

### ^14^C labelled sucrose loading assay

The phloem loading capacity of ^14^C labelled sucrose was quantified following the method as described by Yadav *et al.* (2017). Briefly, rice flag leaves at the milk-stage of grain-filling were cut, and the leaf bases were immediately transferred to petri-dish submerged with MES/CaCl_2_ buffer (20 mM MES, 2mM CaCl_2_, pH 5.5 with KOH). Next, leaf discs (3.6 × 1.0 cm) were excised using a cork borer, and immediately placed abaxial side down in a 24-well microtiter plate pre-filled with 1 ml MES/CaCl_2_ buffer spiked with 1mM sucrose solution (1mM unlabelled sucrose supplemented with 0.81μCi ml^−1^ ^14^C Sucrose). Leaf discs immersed in the above solution were vacuum-infiltrated for at least 20 min with gentle shaking at room temperature. Labelled leaf discs were transferred to a fresh microtiter plate, washed twice with 1.0 ml MES/CaCl_2_ buffer, blot dried on absorbent filter paper, immediately frozen on dry ice, and lyophilized. Next, ^14^C sucrose loading capacity into leaves was quantified by a scintillation counter (PerkinElmer Inc.). Six biological replicates with four leaf discs each were used per genotype for the experiment.

### Sucrose quantification in the phloem sap

The amount of sucrose in the phloem was quantified using the EDTA-facilitated exudation technique described by King and Zeevaart (1974). Flag leaves of the cultivated and wild rice were excised at the milk-stage of grain-filling at around 9-10 am (when leaf photosynthesis was quantified). Leaf bases were immediately recut under exudation buffer (10 mM, Hepes, 10 mM EDTA, pH 7.5) to allow dispersion of any leaf contaminants, and placed in a 15 ml tube containing 5 ml of the exudation buffer. Samples were incubated in a humid chamber (relative humidity > 90%) in the dark to prevent evapotranspiration, and exudates were collected after 5 h of incubation. 2 ml of sample aliquot was spiked with 50 μl internal standard (1 mg ml^−1^ ribitol in water), and freeze-dried for sucrose quantification using GC-MS based method as described earlier. Individual flag leaves from different plants were considered as separate biological replicates, and four biological replicates were analyzed for each genotype.

### RNA isolation, cDNA synthesis, and qRT-PCR analysis

Tissue samples for RNA isolation and qRT-PCR analysis were collected from the same developmental stage of *O. sativa* ssp. *japonica* cv. Nipponbare and *O. australiensis*. Samples were collected at the milk-stage of grain filling (ten days after heading) for the two species from the same regions of flag leaves, stems, panicle bases, and spikelets. Total RNA from flag leaf, stem, and panicle base was extracted using TRIzol reagent (Invitrogen), whereas RNA from spikelet was extracted using plant RNA purification kit (Sigma-Aldrich) according to manufacturer’s protocol. One microgram (1μg) of total RNA was reverse transcribed to first strand-cDNA by anchored oligodT priming using Thermo Scientific RevertAid cDNA synthesis kit following manufacturer’s instructions. The primer pairs (Table S1) of the selected target genes used in this study produced a single product as viewed through dissociation curve analysis. Further, the efficiency of each primer pair was also evaluated using a standard curve method where five cDNA quantities (1, 25, 50, 75, and 100 ng) from each genotype were used to construct a standard curve, and efficiency was calculated using the formula E = (10^[−1/slope]^-1)*100. Expression analysis of each gene was performed in three biological replicates for each genotype. The transcript levels of genes were normalized independently to two stable internal controls actin (LOC_Os03g50885) and ubiquitin (LOC_Os03g03920) with similar results. Relative expressions of target genes are presented using actin as the internal control applying 2^−ΔCt^ method by Livak and Schmittgen (2001). *In silico* expression analysis of the genes was performed using OS_AFFY_RICE_6 dataset present in Genevestigator database (Hruz *et al.*, 2008; https://genevestigator.com/).

### Visualization and quantification of anatomical traits

The vascular features of flag leaves as well as vascular bundles in the inner and outer ring of the stem, and panicle base (0.5 cm above the panicle node) were visualized and quantified using hand-cut transverse sections. Flag leaves, stems, and panicle bases from five independent plants for each genotype were hand-sectioned using a razor-blade at the milk-stage of grain-filling, and stained with toluidine blue O (0.02% toluidine blue O in water) as described by Mitra and Loqué, (2014). Photographs were taken under a bright field microscope (LMI). Quantification of the different anatomical traits was made using Fiji-Image J software (Schindelin *et al.*, 2012).

### Cellulose staining and quantification

For histochemical visualisation of cellulose, transverse hand-sections of rice stem (approximately 20 μm thickness) at the milk-stage of grain-filling were stained using calcofluor white (0.2% calcofluor white M2R in water) staining solution as described by Ambavaram *et al.* (2011). Stained sections were photographed under UV light range using a fluorescence microscope (excitation filter: 340-380 nm; Carl Zeiss Microscope). The quantification of cellulose and hemicellulose was made using National Renewable Energy Laboratory (NREL) protocols as described in Mund *et al.* (2016). The quantifications were made in four biological replications for each genotype.

### Extraction of stem tissues for quantification of sucrose metabolic enzyme activity

The enzyme activity was performed in the stem tissues of *O. sativa* ssp. *japonica* cv. Nipponbare and *O. australiensis*. Tissue sample (100 mg fresh weight) was ground in liquid nitrogen, and homogenized in extraction buffer (50 mM HEPES/NaOH (pH 7.5), 7.5 mM MgCl_2_, 2 mM EDTA, 2% (w/v) PEG 8000, 2% (w/v) PVP and 5 mM DDT) at 4 °C. The homogenate was centrifuged for 1 min at ~16,000 × *g*, the pellet was discarded, and the crude extract was used for enzyme activity assay.

### Estimation of cell wall invertase activity

Cell wall invertase activity was estimated as described in Tomlinson *et al.* (2004). 50 μl of crude enzyme extract was added to 150 μl assay mix, containing 0.1 M sucrose in 50 mM sodium acetate at pH 4.7, on ice. The assay reaction was incubated at 37°C for 30 min. The reaction was alkalinized by the addition of 50 μl 1 M TRIS-HCl (pH 8.0), and then heated at 85°C for 3 min. Two blanks were set up to measure acid hydrolysis of sucrose and endogenous glucose levels. The amount of released hexoses was measured enzymatically using Sucrose/D-Fructose/D-Glucose assay kit (Megazyme), and invertase activity was expressed as described in Nishanth *et al.* (2018).

### Estimation of sucrose synthase synthesis and cleavage activity

To quantify OsSUS synthesis activity, 50 μl of crude extract was incubated with 20 mM HEPES/NaOH buffer (pH 7.5), 5 mM MgCl_2_, 20 mM KCl, 12 mM fructose, 0.4 mM phosphoenolpyruvate, 2 mM uridine diphosphate UDP-glucose, 20 U pyruvate kinase, 20 U lactate dehydrogenase, and 0.15 mM NADH in 1.0 ml final reaction volume. Similarly, to quantify OsSUS cleavage activity, 50 μl of crude extract was incubated with 20 mM HEPES/NaOH buffer (pH 7.5), 100 mM sucrose, 2 mM UDP, 2mM MgCl_2_, 0.005 UDP-glucose dehydrogenase, and 1.5 mM NAD^+^ in 1.0 ml final reaction volume. The reaction mixture was mixed gently, and a decrease and an increase in absorbance were measured for synthesis and cleavage activity, respectively, at 340 nm continuously for 0 to 300 sec using a UV–Vis spectrophotometer. The OsSUS activity was calculated as μmol NADH oxidized (synthesis activity) and NAD+ reduced (cleavage activity) per min per mg protein as described in Qazi *et al.* (2012).

### Subcellular localization of OsSUS1

Full length of *OsSUS1* CDS without stop codon was cloned under constitutive 35S promoter with a C-terminal eYFP fusion into binary vector pEG10 (Earley *et al.*, 2006). The construct along with empty vector control was then, introduced into EHA105 strain of *Agrobacterium tumefaciens*. Each transformant was co-infiltrated with a plasma membrane-localized marker (PM-mCherry; Nelson *et al.*, 2007) into the leaves of *Nicotiana benthamiana*. Confocal laser scanning microscopy TCS SP5 (Leica Microsystems) was used to take the images with appropriate lasers.

## Results

### Biomass and grain yield differences among the selected wild and cultivated rice accessions

We initiated the study with two cultivated rice varieties, *Oryza sativa* ssp. *japonica* cv. Nipponbare and *Oryza sativa* ssp. *indica* cv. IR 64, and three wild relatives of rice, *Oryza rufipogon*, *Oryza latifolia,* and *Oryza australiensis* that show remarkable variations in overall growth and architectural features at both vegetative and reproductive stages (Fig. 1A). All the selected wild rice species grew taller compared to the two cultivated varieties (Fig. 1B). Among all the accessions, *O. sativa* cv. Nipponbare produced the lowest number of total leaves, while *O. rufipogon* showed the highest number of leaves per plant (Fig. 1C). In addition, Nipponbare showed a significantly lower tiller number per plant compared to all the selected accessions (Supplementary Fig. S1B). Selected wild rice species had a larger leaf surface area compared to the cultivated varieties (Fig. 1D). We, then, quantified the number and length of internodes in the main stem, as internodes are important contributors to the biomass of a plant. Cultivated rice varieties had fewer, smaller, and slender internodes in the main stem compared to the wild accessions (Fig. 1E; and Supplementary Fig. S1A). In addition, stem thickness and circumference were also lower for Nipponbare and IR 64 compared to all selected wild species (Supplementary Fig. S1C, D). Thus, plant architectural traits, such as height, branching, and internode features, as well as leaf traits attributed to the high biomass of the selected wild rice. Consistent with this, the selected wild rice also showed higher fresh and dry weight compared to the cultivated varieties. (Fig. 1F, G).

**Fig. 1.**
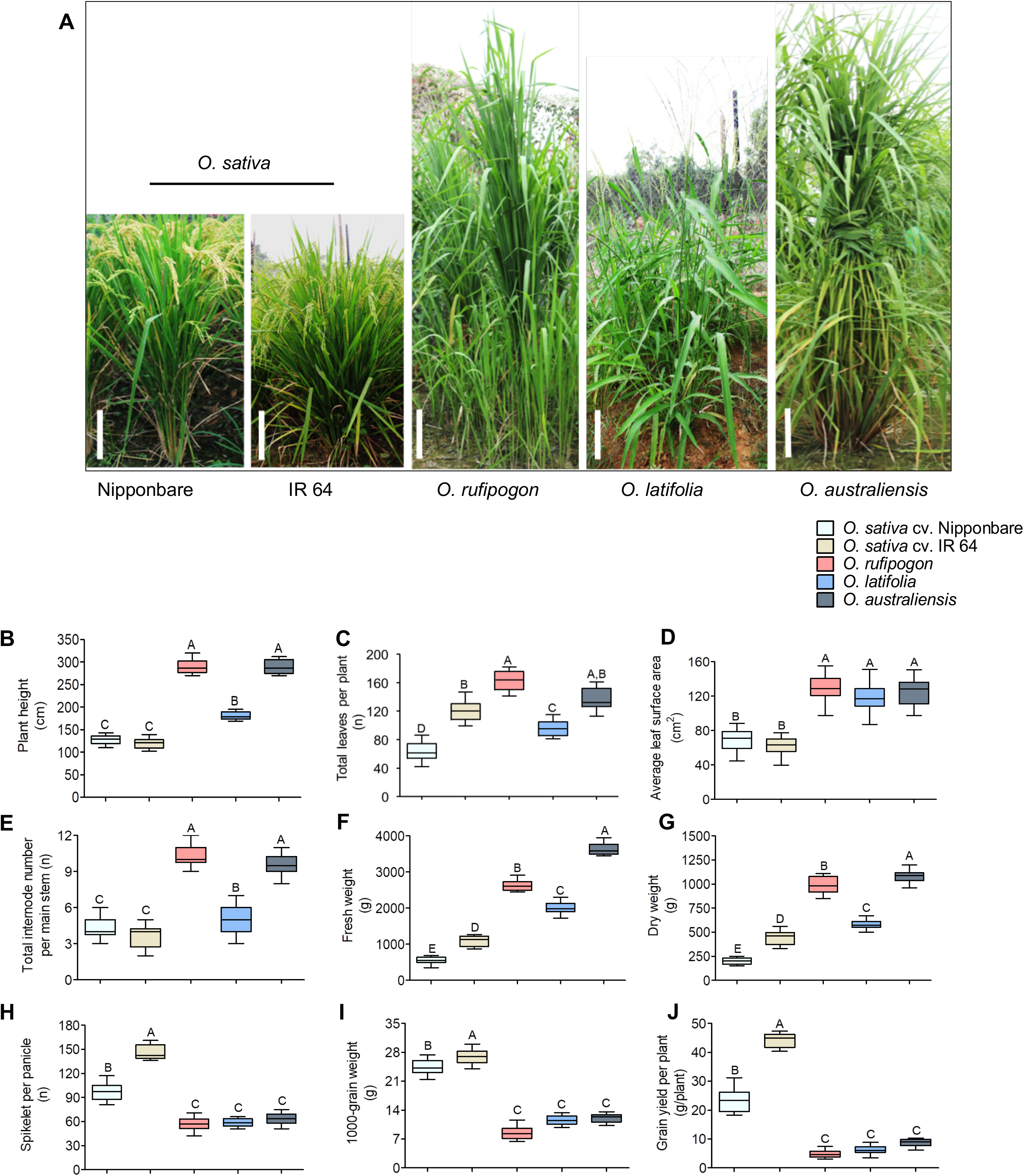
Comparison of biomass and yield-related traits of the selected cultivated and wild rice genotypes. (A) Shown are the photographs of the selected rice genotypes at the reproductive stage (scale bar = 30 cm). (B–J) Quantification of plant height (B), total leaves per plant (C), average leaf surface area (D), total internode number per main stem (E), fresh weight (F), dry weight (G), spikelet per panicle (H), 1000-grain weight (I), and grain yield per plant (J) of the selected rice genotypes. Each box and whisker plot shows the interquartile range with minimum and maximum values of ten data points from different plants. Significance of differences among the genotypes was calculated using One-Way ANOVA with Tukey’s post-hoc test (n = 10, *p* < 0.05).

We, then, quantified seed traits of the selected cultivated and wild rice. The cultivated rice varieties produced higher numbers of spikelets per panicle as well as higher seed weight, resulting in higher grain yield per plant compared to the wild relatives (Fig. 1H-J). Principal component analysis for all the quantified biomass and grain-yield traits showed a clear separation of wild and cultivated genotypes (Supplementary Fig. S2). Taken together, these results confirmed the higher biomass accumulation at the expense of grain yield in the selected wild rice compared to the cultivated varieties. Wild rice *O. australiensis* had higher phenotypic values for all the traits contributing to biomass, including tiller number, compared to Nipponbare. Therefore, we investigated the source strength, sink features, sucrose translocation as well as the fate of the photosynthates towards biomass and yield using representative wild rice and cultivated variety, *O. sativa* cv. Nipponbare and *O. australiensis*, respectively.

### Comparison of source efficiency and photosynthates utilization

In order to compare source efficiency, we quantified leaf photosynthesis rate and related physiological traits as well as soluble sugars/non-structural carbohydrates (NSCs) content in the leaves of the two selected accessions. A significant difference in the leaf photosynthesis rate *A* was observed between the two species at booting, and milk- and dough- stage of grain filling. Both the species showed an increasing trend of leaf photosynthesis from booting to the milk stage, followed by reduction at the dough stage of grain filling (Fig. 2A). The higher photosynthesis in *O. australiensis* at the three stages was also associated with higher *g_s_* and *Ci* compared to Nipponbare (Supplementary Fig. S3A, B). Next, we measured the different soluble sugars/NSCs, such as sucrose, fructose, glucose, and starch, at the time of photosynthesis quantification in leaves at the milk stage of grain filling to evaluate the levels of primary carbon metabolites, the major outcome of photosynthesis. The abundance of all four NSCs was significantly higher in the leaves of *O. australiensis* than Nipponbare (Fig. 2B, C). Interestingly, *O. australiensis* also accumulated higher levels of starch in leaves compared to Nipponbare, suggesting an abundance of residual sugar in the wild rice that is converted to starch. In contrast, the glucose, fructose, and sucrose content as well as starch content were found to be significantly lower in *O. australiensis* grains compared to Nipponbare, despite starch being the major reserve product of seeds (Supplementary Fig. S4). Similarly, seed glucose, fructose, sucrose, and starch content were also found to be significantly lower in two other wild species *O. latifolia* and *O. rufipogon* compared to the cultivated varieties IR 64 and Nipponbare. Higher starch and soluble sugar content in *O. australiensis* leaf and lower starch and soluble sugar levels in seed suggested possible bottlenecks in the mobilization of photosynthates to grains in the wild rice.

**Fig. 2.**
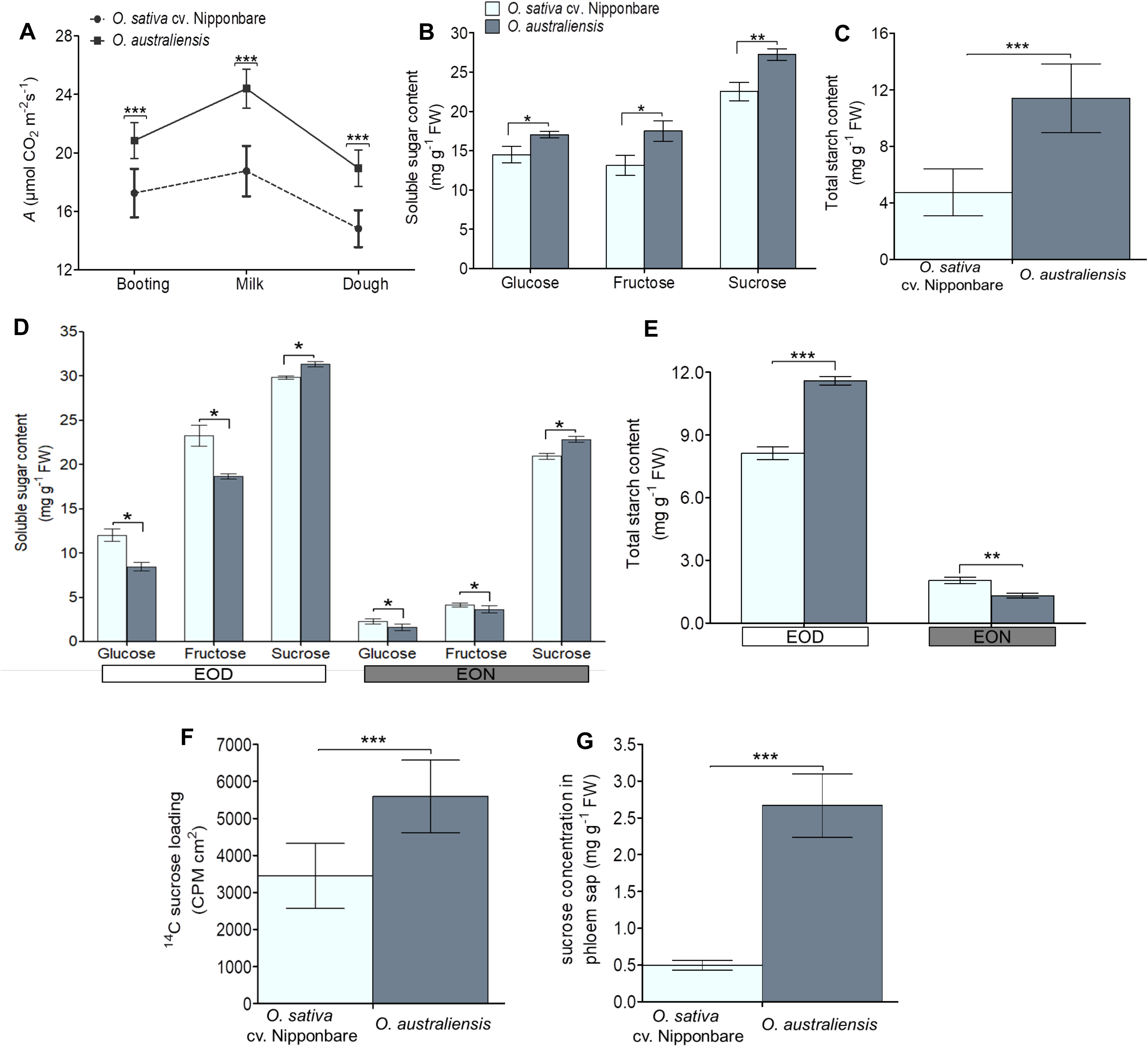
Leaf photosynthesis, sugar accumulation, phloem loading, and sucrose export from leaves of a cultivated rice *O. sativa* cv. Nipponbare and a wild rice *O. australiensis*. (A) Quantification of flag leaf photosynthesis per unit area (*A)* at different stages during booting and grain-filling (n = 15). (B-C) Shown are glucose, fructose, and sucrose content (B), and total starch content (C) in the flag leaves of the cultivated and wild rice at the time of photosynthesis quantification (n = 4). (D-E) Quantification of glucose, fructose, and sucrose content (D), and total starch content (E) in the flag leaves of the cultivated and wild rice at the End Of Day (EOD) and End Of Night (EON) (n = 4). (F) Uptake of [^14^C] sucrose in the flag leaf discs expressed as counts per minute (CPM) per square centimeter of leaf area (n = 6). (G) Sucrose quantification in the phloem sap of the two species (n = 4). Data represent the mean and standard deviation (SD), and significance of differences between the genotypes was calculated using student t-test (*p < 0.05, **p < 0.01, ***p < 0.001).

We, then, quantified end of the day (EOD) and end of the night (EON) carbon status in the leaves of Nipponbare and *O. australiensis*. Glucose and fructose level were significantly lower in *O. australiensis* than *O. sativa* cv. Nipponbare at the EOD and EON (Fig. 2D). Sucrose content was marginally, but significantly, higher in *O. australiensis* compared to *O. sativa* cv. Nipponbare at the EOD as well as EON. At the EOD, *O. australiensis* accumulated more starch in leaves than *O. sativa* cv. Nipponbare (Fig. 2E). However, *O. australiensis* displayed a significantly lower leaf starch level than *O. sativa* cv. Nipponbare at the EON, suggesting efficient utilization of stored starch in *O. australiensis* during the night (Fig. 2E). This is in contrast to lower starch content in the grains of *O. australiensis* as explained earlier (Supplementary Fig. S4). These results indicated the proper utilization of the photosynthates in *O. australiensis*, however not towards grain filling, suggesting the possibility of the alternative sink and alternative fate of photosynthates.

### Phloem loading and export of sucrose from leaves

Since sucrose is the major transportable form of photosynthates from source leaves to sink organs, we investigated the sucrose transport differences between the two accessions. We performed [^14^C] labelled sucrose assay in leaf discs of *O. sativa* cv. Nipponbare and *O. australiensis*. A higher amount of ^14^C per unit leaf disc area was detected in *O. australiensis* (5598.83 CPM per cm^2^) compared to that of in Nipponbare (3446.66 CPM per cm^2^) (Fig. 2F). We, then, quantified sucrose in the phloem sap of the two accessions. Phloem sap of *O. australiensis* had four times higher sucrose content than *O. sativa* cv. Nipponbare (Fig. 2G). Together, ^14^C-assay and analysis of phloem sap showed better phloem loading and sucrose export from the leaf in the wild rice *O. australiensis* compared to Nipponbare.

### Expression pattern of genes encoding sucrose transporters

We examined the contribution of sucrose transporters for differences in phloem loading and sucrose export from leaves. Since clade III SWEET transporters and SUT transporters are the major sucrose transporters in plants, we preferentially quantified the transcript levels of genes encoding those transporters. Expression analysis of *OsSWEET*s using publicly available datasets showed preferentially source-specific expression of *OsSWEET13* (Supplementary Fig. S5A). *OsSWEET14* also followed a similar source-specific expression pattern, but at a relatively lower expression level than *OsSWEET13*. *OsSWEET15* and *OsSWEET11* showed preferentially sink (panicle and endosperm) specific expression patterns. The expression levels of *OsSWEET12* was observed to be very low in all the tissues. Consistent with this, quantitative Real Time-PCR results showed higher transcript levels of *OsSWEET13* in flag leaf and stem; *OsSWEET11* in the spikelet; and *OsSWEET15* in the stem, spikelet, and flag leaf of Nipponbare (Fig. 3A-D). Interestingly, the transcript level of *OsSWEET13*, encoding preferentially source-specific SWEET transporter, was significantly higher in flag leaf and stem of *O. australiensis* compared to Nipponbare (Fig. 3A, B). *In silico* expression analysis showed higher expression of *OsSUT1* and *OsSUT2* in different tissue compared to other *OsSUT*s (Supplementary Fig. S5B). qRT-PCR analysis confirmed the generally higher expression of *OsSUT1* and *OsSUT2* in different tissues of Nipponbare (Fig. 3E-H). We detected higher transcript levels of *OsSUT1* in the flag leaf, and *OsSUT2* in the stem of *O. australiensis* compared to Nipponbare (Fig. 3E, F). The expression pattern of *OsSWEET*s and *OsSUT*s suggested the potential involvement of selected transporters, such as OsSWEET13, OsSUT1, and OsSUT2, for higher phloem loading and sucrose export from leaf to stem in the wild rice.

**Fig. 3.**
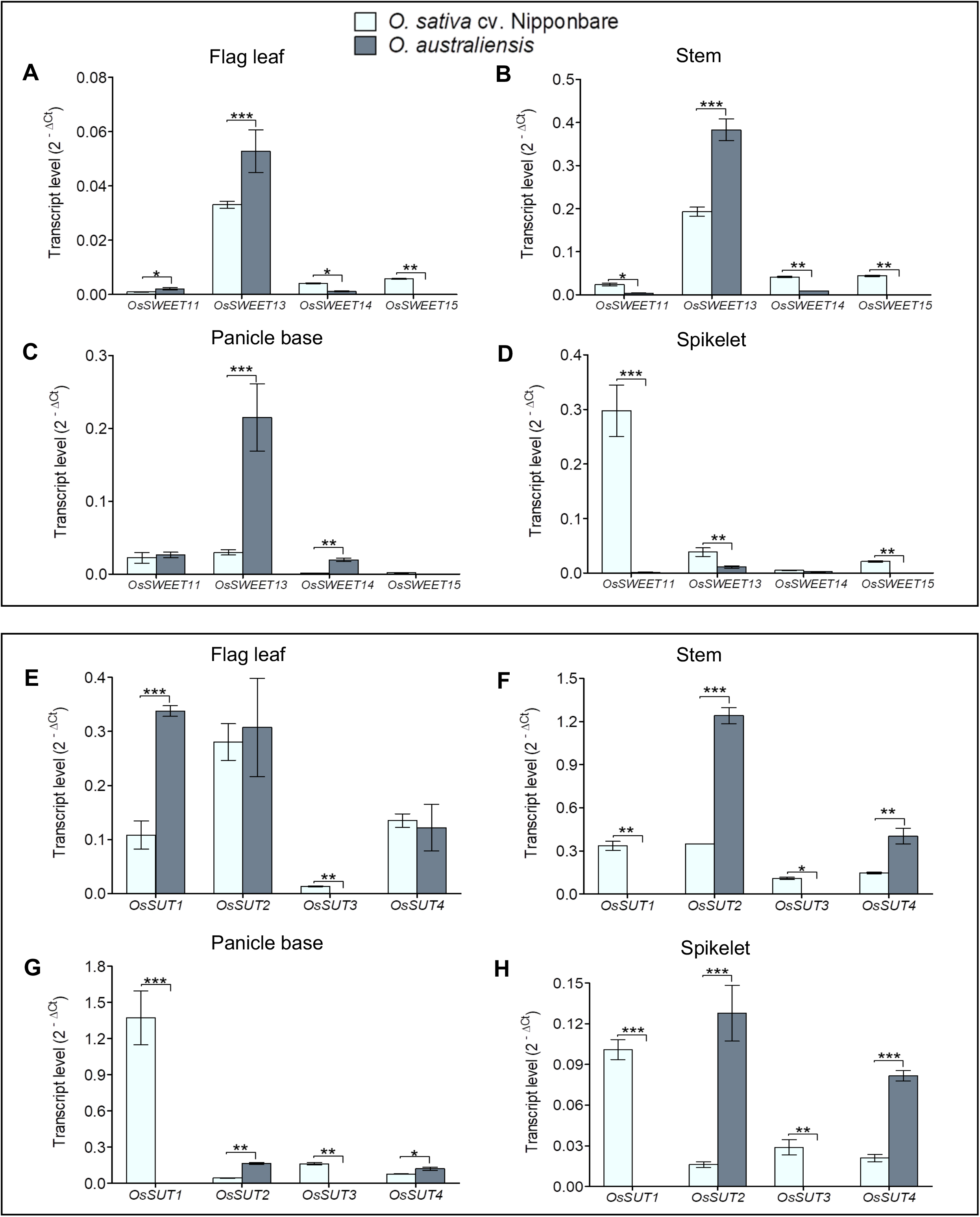
Tissue-specific expression pattern of genes encoding SWEET and SUT transporters in a cultivated rice *O. sativa* cv. Nipponbare and a wild rice *O. australiensis*. (A–D) Transcript levels of *SWEET* genes in the flag leaf (A), stem (B), panicle base (C), and spikelet (D) at the milk-stage of grain-filling in the two rice species. (E–H) Transcript levels of *SUT* genes in the flag leaf (E), stem (F), panicle base (G), and spikelet (H) at the milk-stage of grain-filling in the two rice species. Data represent the mean (n = 3) and standard deviation (SD), and significance of differences between the genotypes was calculated using student t-test (*p < 0.05, **p < 0.01, ***p < 0.001).

Despite higher leaf photosynthesis rate and more sucrose export from source leaves, *O. australiensis* produced smaller and lighter grains with less starch content compared to the cultivated variety Nipponbare. Therefore, we also examined the transcript levels of different sucrose transporters in developing spikelets. The expression levels of all the *OsSWEET*s were significantly lower in spikelets of *O. australiensis* compared to Nipponbare, with almost no expression of preferentially sink-specific *OsSWEET15* and *OsSWEET11* (Fig. 3D). In addition, the transcript levels of *OsSUT1* was significantly lower in developing spikelet of *O. australiensis* than Nipponbare (Fig. 3H). Thus, reduced grain filling in *O. australiensis* could potentially be associated with reduced expression of *OsSWEET15*, *OsSWEET11*, and *OsSUT1*.

### Differences in vascular features associated with sucrose transport

Vascular features are also key to photoassimilate partitioning from source to sink tissues. Therefore, we quantified the vein number and vein width in the fully matured flag leaves of *O. sativa* cv. Nipponbare and *O. australiensis*. *O. australiensis* leaves had a higher number of wider veins compared to Nipponbare (Fig. 4A, B). Moreover, *O. australiensis* also exhibited higher vein density with lesser interveinal distance compared to Nipponbare (Supplementary Fig. S6A, B). We, then, checked the organization of vascular bundles at the panicle base of the two accessions, as the panicle base is the site of attachment of spikelets to the main plant (Zhang *et al.,* 2002; Zhai *et al.,* 2018). *O. australiensis* had a fewer number of vascular bundles with a reduced area than Nipponbare at the panicle base (Fig. 4C, D). We also quantified the vascular features in the stem of the two accessions (Fig. 4E, F). Interestingly, we observed ~1.7 times larger vascular bundles in the *O. australiensis* stem compared to Nipponbare. Further, *O. australiensis* stem also had approximately twice the number of vascular bundles than Nipponbare stem due to wider stem (Fig. 4F). We also checked the vascular features in additional cultivated and wild rice accessions, *O. sativa* cv. IR 64 and *O. latifolia*, respectively. The results were found to be consistent with the differences between Nipponbare and *O. australiensis* for flag leaves, stem, as well as panicle base (Supplementary Fig. S7). Taken together, the sucrose mobilization to a panicle in *O. australiensis* might likely be hindered due to defects in the vascular bundle at the panicle base and minimal expression of *OsSWEET*s in developing spikelets, strengthening the hypothesis of alternative utilization of photosynthates in the *O. australiensis* stem towards higher biomass.

**Fig. 4.**
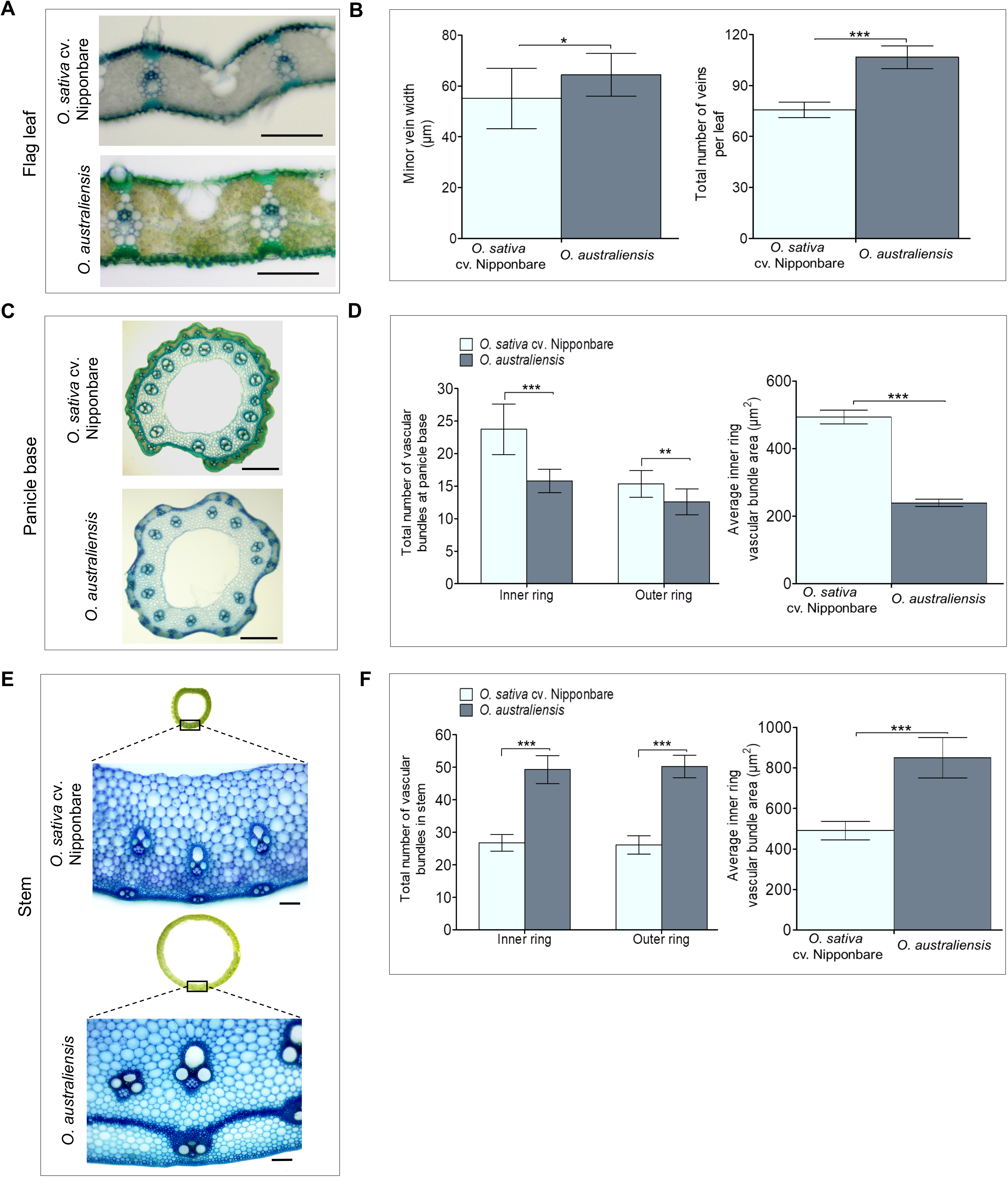
Vascular features in flag leaf, panicle base, and stem of a cultivated rice *O. sativa* cv. Nipponbare and a wild rice *O. australiensis*. (A-B) Cross-sections of flag leaves of the two species (A, scale bar = 100 μm), and quantification of minor vein width and the total number of veins (B). (C-D) Transverse sections at the panicle base (0.5 cm above panicle node) of the two species (C, scale bar = 250 μm), and quantification of number and area of vascular bundles (D). (E-F) Transverse sections of stems of the two species (E, scale bar = 100 μm), and quantification of number and area of vascular bundles (F). The peripheral concentric ring of the panicle base and stem is represented as the outer ring, and the inner wider circle is represented as the inner ring in figures. Data represent the mean and standard deviation (SD) from fifteen cross-section images collected from five different plants (n = 5), and significance of differences between the genotypes was calculated using student t-test (*p < 0.05, **p < 0.01, ***p < 0.001).

### Starch accumulation and expression levels of starch-biosynthesis genes in stem

The higher leaf photosynthesis rate and phloem loading in leaves, along with vascular features and the expression pattern of sucrose transporter genes indicated stem to be a major sink organ utilizing sucrose received from leaves in *O. australiensis*. Therefore, we quantified the content of NSCs in stems of the two species at the milk-stage of grain filling (Fig. 5A, B). Significantly lower amounts of glucose, fructose, and sucrose were detected in the stem of *O. australiensis* as compared to Nipponbare (Fig. 5A). This was in contrast to a higher amount of sucrose being exported from leaves of *O. australiensis*. Therefore, we suspected that *O. australiensis* might reserve a high amount of starch in their stem. To our surprise, a negligible amount of starch was present in the stem of *O. australiensis* compared to Nipponbare (Fig. 5B). Starch content was, further, found to be very low in the *O. australiensis* stems at different stages and in different internodes (Fig. 5C, D). Starch content in the stems of another cultivated variety, *O. sativa* cv. IR 64, and wild species, *O. latifolia* and *O. rufipogon*, also showed a similar pattern. Similar to *O. australiensis*, a negligible amount of starch was present in the stems of *O. latifolia* and *O. rufipogon* compared to IR 64 (Supplementary Fig. 8). We, then, checked the expression of genes involved in starch biosynthesis. Since we were hypothesizing differential fate of sucrose in the stem, we initially checked the expression of members of ADP-glucose pyrophosphorylase large subunit (*OsAPL*) and small subunit (*OsAPS*), and starch synthase (*OsSS*) gene families in internode, node, and culm of Nipponbare using publicly available data at Genevestigator (Hruz *et al.*, 2008; https://genevestigator.com/). We observed generally high expression of these genes in the internode, node, and stem of the cultivated variety (Supplementary Fig. 9A, B). Since *OsAPL3*, *OsAPS1,* and *OsSSIIb* from the starch biosynthesis gene families showed the highest expression levels in the internode, we selected these genes for expression analysis at multiple tissues using Genevestigator as well as qRT-PCR validation in the stem. Expectedly, very high expressions of these genes were detected in the internode and stem of the cultivated variety Nipponbare by both *in silico* expression analysis as well as qRT-PCR analysis (Fig. 5E; Supplementary Fig. S11A). Consistent with very low starch levels in *O. australiensis* stem and internode, the transcript abundance of genes encoding two key enzymes, *OsAPL3* and *OsAPS1*, was drastically less in *O. australiensis* compared to Nipponbare (Fig. 5E). A lower amount of soluble non-structural carbohydrates, in particular remarkably low starch content, despite more sucrose transport from leaves to stem, confirmed an alternative fate to photosynthates in *O. australiensis* stem.

**Fig. 5.**
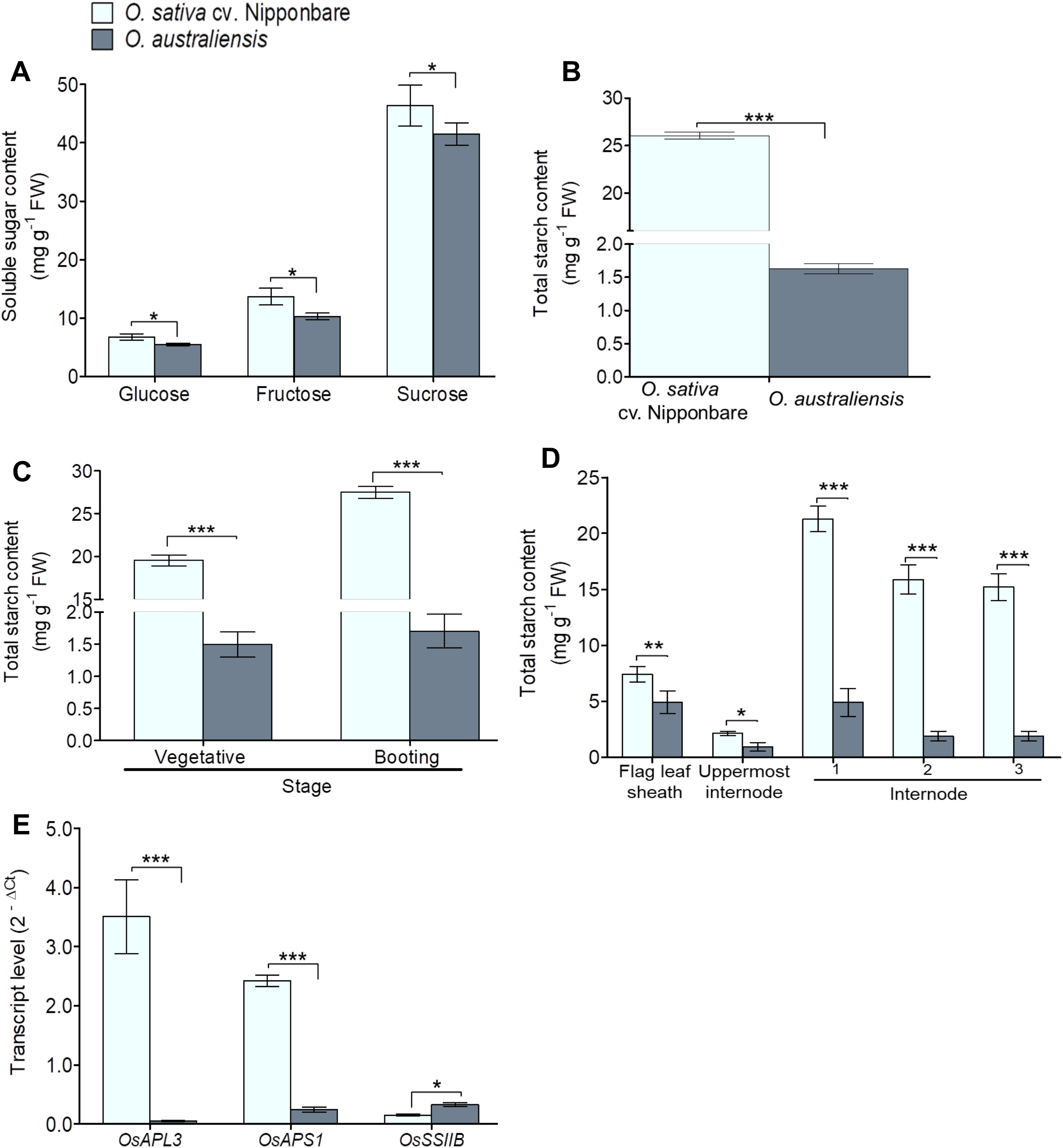
Quantification of soluble sugars and expression pattern of starch-biosynthesis genes in stems of a cultivated rice *O. sativa* cv. Nipponbare and a wild rice *O. australiensis*. (A-B) Glucose, fructose, and sucrose content (A), and total starch content (B) in the stem of the two species at the milk-stage of grain-filling (n = 4). (C) Quantification of total starch in the stem at the vegetative and booting stage of the two species (n = 4). (D) Starch content in the flag leaf sheath and different internodes at milk-stage of grain-filling of the two species (n = 4). (E) Transcript levels of starch-biosynthesis genes, *OsAPL3* (encoding ADP-glucose pyrophosphorylase large subunit), *OsAPS1* (encoding ADP-glucose pyrophosphorylase small subunit), and *OsSSIIB* (encoding Starch synthase) in the stem of the two species at milk-stage of grain-filling (n = 3). Data represent the mean and standard deviation (SD), and significance of differences between the genotype was calculated using student t-test (*p < 0.05, **p < 0.01, ***p < 0.001).

### Expression levels of sucrose metabolism genes, sucrose synthase and cell wall invertase enzyme activity, and structural carbohydrate levels in stem

Invertases (INV) and sucrose synthase (SUS) are the key enzymes for the degradation of sucrose. Expression of genes encoding invertases has been shown to be strongly correlated with starch synthesis (Bahaji *et al.*, 2014; Ruan, 2014). An *in silico* expression analysis of members of Invertase gene-family, including cell wall invertase (*OsCIN*), cytoplasmic invertase (*OsNIN*), and vacuolar invertase (*OsINV*), showed high expression of many of those genes in the stem, node, and internode tissues of the cultivated rice (Supplementary Fig. S10A). Previously, *OsCIN1* and *OsINV2* were shown to be highly abundant in rice stem as compared to other members of the gene-family (Ji *et al.*, 2005). We, then, checked the expression pattern of representative genes encoding cell wall invertase (*OsCIN1*), cytoplasmic invertase (*OsNIN8*), and vacuolar invertase (*OsINV2*) in stem tissues by qRT-PCR (Fig. 6A). The expression of *OsCIN1* was drastically lower in *O. australiensis* stem compared to Nipponbare. In addition, *O. australiensis* also showed lower cell wall invertase activity in the stem (Fig. 6B). Together, limited expression of a cell wall invertase gene along with lower cell wall invertase enzyme activity would limit the breakdown of sucrose in the *O. australiensis* stem.

**Fig. 6.**
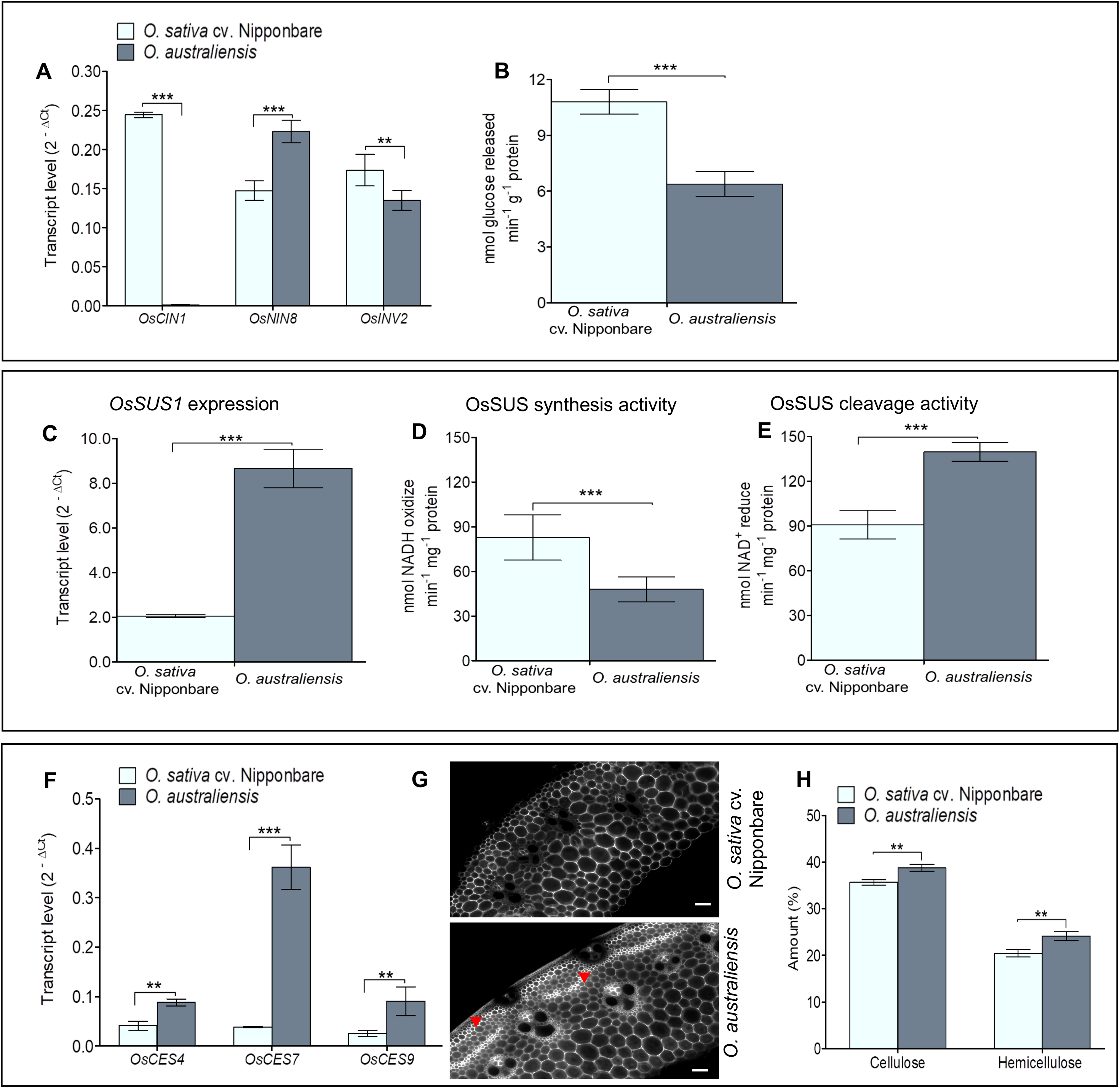
Gene expression and activities of sugar metabolic enzymes, and structural carbohydrates levels in stem tissue of a cultivated rice *O. sativa* cv. Nipponbare and a wild rice *O. australiensis* at milk-stage of grain-filling. (A) Transcript levels of genes encoding cell wall invertase (*OsCIN1*), cytoplasmic invertase (*OsNIN8*), and vacuolar invertase (*OsINV2*) (n = 3). (B) Cell wall invertase enzyme activity (n = 4). (C) Transcript levels of *Sucrose Synthase* 1 (*OsSUS1*) (n = 3). (D-E) OsSUS enzyme activity in synthesis (D) and cleavage (E) direction (n = 4). (F) Transcript levels of *Cellulose Synthase* genes, *OsCES4*, *OsCES7*, and *OsCES9* (n = 3). (G) Calcofluor-white staining for cellulose deposition in the transverse stem sections of the two species. Red arrowhead shows higher deposition of cellulose in *O. australiensis* stem. Scale bar represents 500 μm. (H) Quantification of cellulose and hemicellulose in the stems of the two species (n = 4). Data represent the mean and standard deviation (SD), and significance of differences between the genotypes was calculated using student t-test (*p < 0.05, **p < 0.01, ***p < 0.001).

Among all the *OsSUS*s, *OsSUS1* was shown to be highly expressed in internodes (Hirose *et al.*, 2008). *In silico* expression analysis also showed high expression of *OsSUS1* in internode, node, and culm (Supplementary Fig. S10B, S11A). Therefore, we checked the intracellular localization of OsSUS1 and the expression levels of the corresponding gene. The OsSUS1-YFP signal overlapped with the plasma membrane-localized marker (PM-mCherry), confirming OsSUS1 localization in the plasma membrane (Supplementary Fig. S12). *OsSUS1* was expressed four times higher in *O. australiensis* stem than Nipponbare, indicating a possibility of cellulose accumulation in *O. australiensis* stem (Fig. 6C). Since SUS catalyzes a reversible cleavage of sucrose, it has a synthesis activity promoting sucrose synthesis and a cleavage activity promoting sucrose degradation. The enzyme activity assay revealed a higher OsSUS1 synthesis activity in Nipponbare, while higher cleavage activity in *O. australiensis* (Fig. 6D, E). Lesser accumulation of starch, more transcript levels of *OsSUS1*, and higher cleavage activity of OsSUS1 in *O. australiensis* stem compared to Nipponbare along with localization of OsSUS1 to plasma membrane prompted us to check the cellulose content in the stems of the two accessions. Cellulose synthase genes are usually expressed at the highest levels in rice internodes compared to other tissues as evident from *in silico* expression analysis (Supplementary Fig. S10C, S11B). Interestingly, transcript levels of *OsCES4*, *OsCES7*, and *OsCES9*, key cellulose synthesis genes, were significantly higher in the stems of *O. australiensis* than in Nipponbare (Fig. 6F). Calcofluor-white staining along with cellulose quantification confirmed more cellulose deposition in *O. australiensis* stem compared to Nipponbare (Fig. 6G, H). Like cellulose, hemicellulose content was also significantly higher in *O. australiensis* stem compared to Nipponbare (Fig. 6H). We also compared cultivated rice *O. sativa* cv. IR 64 and wild rice *O. latifolia*, and observed more cellulose deposition in *O. latifolia* stem compared to IR 64 (Supplementary Fig. S13). Taken together, lower expression and activity of cell wall invertase, higher expression and cleavage activity of OsSUS1 coupled with higher expression of cellulose synthase genes, at least in part, led to the utilization of photosynthates in the *O. australiensis* stem towards cellulose deposition.

## Discussion

We investigated the differences in source-sink dynamics between a cultivated rice variety *O. sativa* cv. Nipponbare, which is optimized for high grain yield, and a wild relative of rice *O. australiensis*, which accumulates high biomass with poor grain yield. *O. australiensis* had higher source strength, as evident by consistently higher leaf photosynthesis rate compared to Nipponbare, at least in part due to vascular features of the leaf (Fig. 2A and Fig. 4A, B). High leaf photosynthesis, usually, results in high biomass or yield depending upon preferential sink tissues, provided efficient transport of photoassimilates from leaves (Burnett *et al.*, 2016; Fabre *et al.*, 2020; Fernie *et al.*, 2020). Limitations in sucrose export promote accumulation of sugars in leaves, thereby inhibiting leaf photosynthesis. Therefore, an efficient sucrose export system from leaf would be warranted for the realization of higher source strength of the wild rice to high biomass and/or yield. ^14^C sucrose uptake assay, as well as sucrose content in phloem sap, confirmed a better sucrose export system from wild rice *O. australiensis* leaves (Fig. 2F, G). Larger vascular bundles and fewer number of mesophyll cells between two consecutive veins potentially led to the export of more sucrose from leaves in *O. australiensis*, as reported in different plant systems (Qi *et al.*, 2008; Fujita *et al.*, 2013; Mathan *et al.*, 2016). In addition, higher expression of genes encoding sucrose transporters, OsSWEET13 and OsSUT1, would also facilitate the phloem loading in the wild rice (Fig. 3). High expression of *SbSWEET8-1* of sorghum, a close homolog of *OsSWEET13*, in leaf and its function in phloem loading supports our idea of the important role of *OsSWEET13* in phloem loading (Mizuno *et al.,* 2016). Similarly, *OsSUT1* not only functions for enhancing phloem loading for sucrose transport but also for retrieval of sucrose from the apoplasm along the transport pathway (Scofield *et al.*, 2007). Taken together, high photosynthesis per unit leaf area coupled with efficient export of photosynthates from the leaves of *O. australiensis* suggested that the wild rice had a higher source strength than Nipponbare. A higher amount of soluble sugars in *O. australiensis* leaves compared to the cultivated variety, further, supported the higher source strength of the wild rice (Fig. 2 B, C).

The better grain filling and higher grain yield of the cultivated rice Nipponbare, despite the relatively lower source strength compared to *O. australiensis*, could be explained by the higher number of larger vascular bundles at the panicle base as well as increased expression of relevant sucrose transporter genes at the panicle base and spikelets of the cultivated rice (Fig. 3, 4). Panicle architecture, a key determinant of rice grain yield, is reported to be shaped by the vascular pattern (Sasahara *et al.*, 1999; Terao *et al.*, 2010). Reduced expression of *OsSWEET13* and *OsSWEET15* in the developing spikelet, and of *OsSUT1* at the panicle base and developing spikelet, along with a lower number of smaller vascular bundles at the panicle base limited mobilization of photosynthates to grains in *O. australiensis* compared to the cultivated rice. In agreement with this, *ossweet11 ossweet15* double mutant has been reported to show a smaller seed size (Yang *et al.*, 2018). Similarly, RNA antisense lines of *OsSUT1* showed reduced grain filling and grain weight (Scofield *et al.*, 2007). Increased supply of larger vascular bundles to rice rachis has been shown to promote a higher number of grains per panicle (Zhang *et al.*, 2002; Terao *et al.*, 2010; Zhai *et al.*, 2018). Altogether, higher source strength and limited grain sink strength in *O. australiensis*, attributed to vascular features and sucrose transporter functions, indicated preferential utilization of photosynthates in stem and internodes, contributing to source-sink relationship differences between the selected cultivated and wild rice. In conjunction with this, larger vascular bundles would facilitate efficient sucrose transport and unloading in the wild rice stem. Such vascular features in the stem would, in turn, also provide mechanical strength to support the high biomass of *O. australiensis* (Aohara *et al.*, 2009). Our expression results indicated the potential involvement of OsSWEET13 and OsSUT2 in unloading a larger amount of sucrose in the stem of the wild rice.

There is a competition among the sink tissues for utilization of photosynthates (Patrick *et al.*, 2013; Durand *et al.*, 2018). According to the EcoMeristem model, the final plant architecture is an outcome of competition for resources among different plant parts that depends on photoassimilate-partitioning patterns (Luquet *et al.*, 2006). The higher number of longer internodes and thicker stem with larger leaves, driving high biomass in *O. australiensis* compared to the cultivated variety, suggested higher utilization of photosynthates for vegetative growth. The higher biomass and larger organ size of the wild rice would require more photosynthates for general respiration and maintenance. Efficient sucrose export from leaves, as shown by higher ^14^C phloem loading and sucrose content in the phloem sap, would fulfill the higher requirement of photosynthates in the wild rice. Sucrose quantification in the phloem exudates showed a remarkably higher amount of sucrose in the phloem sap of *O. australiensis* compared to the cultivated variety, whereas the ^14^C sucrose uptake assay showed ~1.6-times higher phloem loading in the wild rice. The artifact-prone nature of the exudation experiments and/or larger leaf area of the wild rice might explain the discrepancy between the sucrose uptake assay and the phloem exudate analysis (Xu *et al.*, 2018). Nonetheless, a higher amount of sucrose loaded into the phloem and exported from the leaves to stem would support the larger vegetative organ size and number, and the general maintenance of the higher biomass in the wild rice.

The preferential utilization of photosynthates in the stem/internodes of wild rice *O. australiensis* in contrast to grains in the cultivated rice Nipponbare suggested differences in the photosynthates metabolism in the stem of the two species. Cultivated rice variety clearly accumulated higher starch content in the stem, which is mobilized to panicle during grain filling (Fig. 5B). *OsSWEET11* and *OsSWEET15* have been suggested to be important for the remobilization of carbon reserve from stem to grain, and OsSUT1 in the retrieval of sucrose from apoplasmic space to stem for conversion to transitory starch (Wang *et al.*, 2020). Indeed, the Nipponbare stem showed higher expression of *OsSWEET11*, *OsSWEET15*, and *OsSUT1* compared to the wild rice along with desirable vascular features at the panicle base. High expression of starch biosynthesis genes, further, contributed to the roles of the stem as an effective source in the cultivated variety at the grain filling stage (Fig. 7). The differences in the stem starch content between the two genotypes projected the stored transitory starch in the stem as a key for source-sink dynamics favoring high grain yield.

**Fig. 7.**
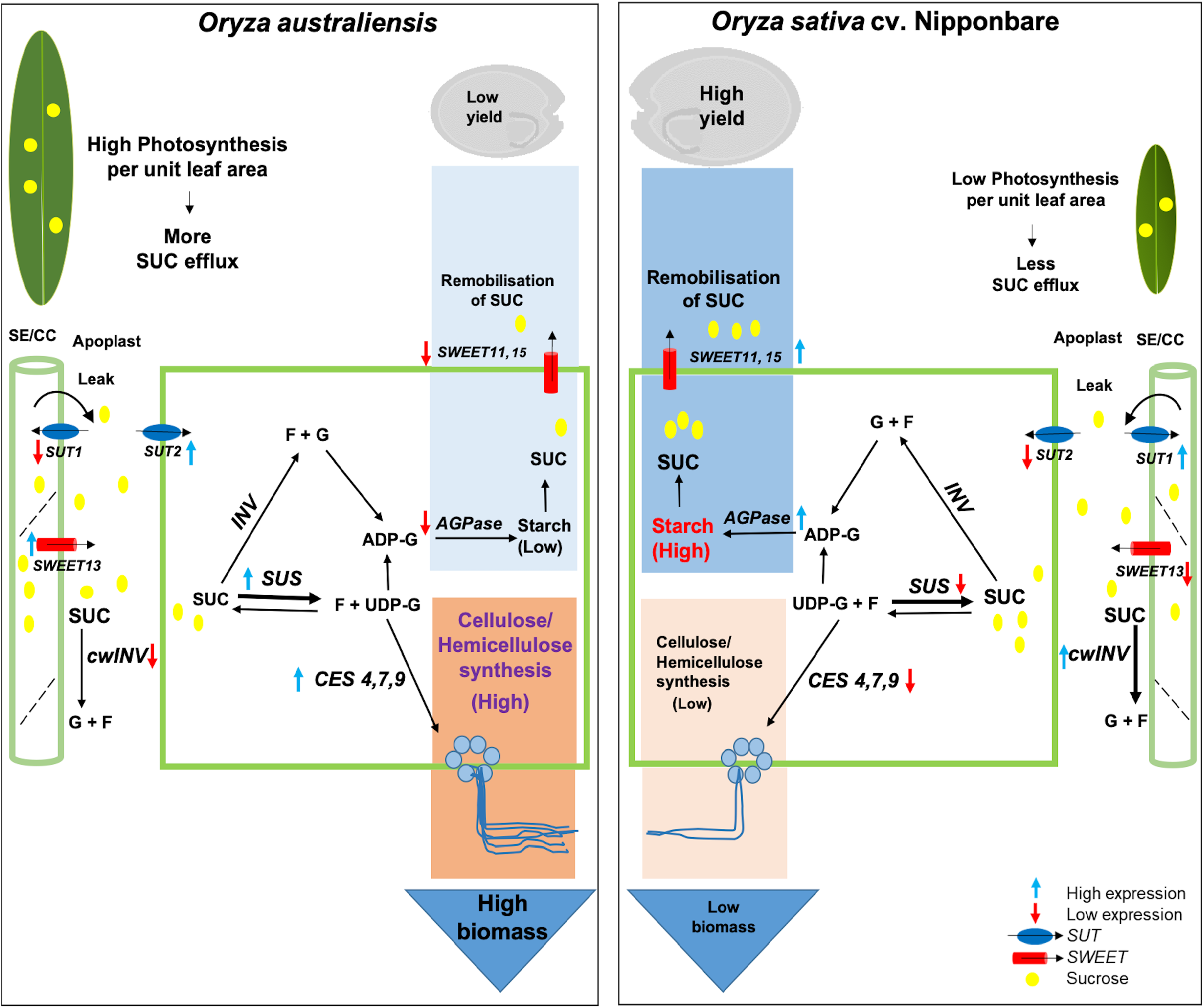
A model explaining the contribution of sucrose transport, sucrose metabolic enzyme activity, starch biosynthesis, and synthesis of structural carbohydrates towards yield and biomass differences between a wild rice *O. australiensis* and a cultivated rice *O. sativa* cv. Nipponbare. High leaf photosynthesis rate coupled with higher expression of selected *OsSWEET* and *OsSUT* genes mediate export of a higher amount of sucrose from a leaf in *O. australiensis* compared to *O. sativa* cv. Nipponbare. A higher amount of sucrose gets unloaded into the stem of *O. australiensis* due to the functions of *OsSWEET13* and *OsSUT2*. However, a lower expression of a gene encoding cell wall invertase (OsCIN1) along with the lower activity of cell wall invertase would limit the formation of glucose and fructose in the wild rice stem. Sucrose gets converted to UDP-G more efficiently in the wild rice *O. australiensis* due to higher expression of *OsSUS1* and more cleavage activity of OsSUS compared to the cultivated rice Nipponbare. Higher expressions of *OsCES4, OsCES7,* and *OsCES9*, then, promote the synthesis of cellulose in the stem of the wild rice. In contrast, higher expressions of starch-biosynthesis genes (*AGPase*, small and large subunit) lead to higher starch content in the stem of the cultivated rice. The higher synthesis activity of OsSUS along with the higher expression of *OsSWEET11* and *OsSWEET15* in the cultivated rice would facilitate efficient remobilisation of sucrose from stem to panicles at the grain-filling stage. SUC, sucrose (yellow colour circle); F, fructose; G, glucose; UDP-G, UDP-glucose; ADP-G, ADP-glucose; *INV*, invertase; cwINV, cell wall invertase; SUS, sucrose synthase.

Cell wall invertases mediate the breakdown of sucrose into glucose and fructose, which enter the cytoplasm through H+/hexose symporters (HXTs) (Ruan *et al.*, 2010). Lower expression and activity of cell wall invertase in the *O. australiensis* stem, which receives a higher amount of sucrose from the leaves, would limit the formation of glucose and fructose. In contrast, higher expression and activity of cell wall invertase in Nipponbare would generate relatively more hexoses that may facilitate the hexose transport pathway to promote starch biosynthesis. The cleavage activity of SUS enzyme would produce UDP-glucose from sucrose in the *O. australiensis* stem, which eventually would promote biomass accumulation through cellulose deposition (Stein and Granot, 2019). Consistent with this, *O. australiensis* stem showed a higher expression of *OsSUS1* and higher cleavage activity SUS compared to Nipponbare. In addition to serving as the prime source for cellulose synthesis, UDP-glucose can also be converted to starch (Asano *et al*., 2002; Koch, 2004; Smith *et al*., 2012)). However, the possibility was minimized in *O. australiensis* due to the very limited expression of *OsAPL3* and *OsAPS1*. Thus, most of the SUS-generated UDP-glucose was converted to cellulose, supported by high expression of cellulose synthase genes, *OsCES4*, *OsCES7*, and *OsCES9* in *O. australiensis* stem (Fig. 6F). OsSUS1 has been reported to be located in SE-CC complexes of phloem, and localization of SUS in phloem is important for cellulose synthesis (Smith *et al*., 2012; Regmi *et al.*, 2016). Plasma membrane localization of OsSUS1 in this study together with earlier reported association of cellulose synthase complexes to plasma membrane highlighted the key importance of OsSUS1 for cellulose synthesis in rice (Supplementary Fig. S12, Lei *et al*., 2012). In addition, SUS functions in companion cells of phloem have been suggested to enhance the sucrose unloading at sink tissues, further helping in unloading photosynthates in the wild rice stem (Nolte and Koch, 1993; Stein and Granot, 2019, Yao *et al.*, 2020). The potential role of SUS in cellulose synthesis, as well as in secondary cell wall thickening, has been investigated in the past (Coleman *et al.*, 2009; Baroja-Fernández *et al.*, 2012; Wei *et al.*, 2015). Consistent with the higher expression of *OsSUS1* in *O. australiensis*, overexpression of *SUS* has been shown to result in increased vegetative growth rate, plant height, and biomass in different plant species (Coleman *et al.*, 2006; Nguyen *et al.*, 2016; Stein and Granot, 2019). Contrary to *O. australiensis*, higher synthesis activity of SUS in the Nipponbare stem might contribute to the sucrose synthesis to be mobilized for grain filling at the heading stage. The differences between the synthesis and cleavage activities of SUS between the two species could be due to many potential factors, such as pH and metabolic status of the tissue, cellular localization, phosphorylation, and oligomerization status of the enzyme (Schmolzer *et al.*, 2016; Stein and Granot, 2019; Takeda *et al.*, 2017). Our results on differential activity of cell wall invertase and starch levels in the stem suggested that the metabolic status of the stem could likely be an important determinant for the differences in synthesis and cleavage activity of SUS between the two species. Further extensive biochemical investigations would be required to establish the role of the metabolic status as well as other contributing factors for the observed differences in SUS synthesis and cleavage activity. Nonetheless, differential functions of cell wall invertase and sucrose synthase between *O. australiensis* and Nipponbare would, at least in part, explained the differential fate of photosynthates in the stem (Fig. 7).

In summary, differences in vascular features and sucrose transporter functions led to a differential source-sink relationship between wild rice *O. australiensis* and cultivated variety *O. sativa* cv. *Nipponbare*. *O. australiensis* showed source-sink dynamics favoring high biomass through the accumulation of structural carbohydrates, mediated by lower cell wall invertase activity, higher SUS cleavage activity together with higher expression of genes encoding cellulose synthases (Fig. 7). In contrast, source-sink dynamics favored higher grain yield in Nipponbare via accumulation of transitory starch in the stem, to be mobilized to panicles with the onset of grain filling. Taken together, vascular features and sucrose transporter functions along with transitory starch storage mechanism and invertase and SUS enzyme activity can potentially be targeted for source-sink dynamics favoring either biomass accumulation in fodder crops or higher grain yield in cereal crops.

## Supplementary data

Fig. S1. Internode length, tiller number, and stem features of the selected cultivated and wild rice genotypes.

Fig. S2. Principal component analysis of biomass and yield traits of the selected cultivated and wild rice species.

Fig. S3. Stomatal conductance (*g_s_*) and intercellular CO2 concentration (*C_i_*) of a cultivated rice *O. sativa* cv. Nipponbare and a wild rice *O. australiensis* at different stages during booting and grain-filling.

Fig. S4. Quantification of soluble sugars in matured seeds of the selected cultivated and wild rice genotypes.

Fig. S5. *In silico* organ-specific expression analysis of rice genes encoding clade III *SWEET* and *SUT* transporters using publicly available data at Genevestigator (https://genevestigator.com/).

Fig. S6. Quantification of leaf interveinal distance and vein density for a cultivated rice *O. sativa* cv. Nipponbare and a wild rice *O. australiensis*.

Fig. S7. Vascular features in flag leaf, panicle base, and stem of a cultivated rice *O. sativa* cv. IR 64 and a wild rice *O. latifolia*.

Fig. S8. Total starch content in the stem of a cultivated rice *O. sativa* cv. IR 64 and two wild rice species, *O. rufipogon* and *O. latifolia*.

Fig. S9. *In silico* expression analysis of rice starch biosynthesis genes using publicly available data at Genevestigator (https://genevestigator.com/) in culm (stem), node, and internode.

Fig. S10. *In silico* expression analysis of genes encoding rice invertases (A), sucrose synthases (B), and cellulose synthases (C) in culm (stem), node, and internode using publicly available data at Genevestigator (https://genevestigator.com/).

Fig. S11. *In silico* expression pattern of the selected key genes involved in sugar metabolism in rice using Genevestigator database (https://genevestigator.com/) across multiple different tissues.

Fig. S12. Localization of OsSUS1-YFP in the plasma membrane of leaf epidermal cells of *Nicotiana benthamiana*.

Fig. S13. Calcofluor-white staining for cellulose deposition in the transverse stem sections of a cultivated rice *O. sativa* cv. IR 64 and a wild rice *O. latifolia*.

Table S1. List of primer pairs used in the study

## Acknowledgements

This work was supported by the Innovative Young Biotechnologist Award (BT/09/IYBA/2015/01) and Ramalingaswamy Re-entry Fellowship (BT/RLF/reentry/05/2013) to AR from the Department of Biotechnology, Ministry of Science and Technology, India. JM and AS acknowledge their CSIR-JRF and SERB-NPDF fellowships, respectively. We also acknowledge Central Instrument Facility, NIPGR; Advanced Instrumentation Research Facility, JNU; Confocal Microscopy Facility, NIPGR; and DBT-eLibrary Consortium (DeLCON) for providing access to e-resources.

## Author Contributions

JM and AR conceptualized the study and designed experiments. JM performed experiments. JM and AS analysed data. JM, AS and AR wrote the manuscript. All the authors have read and edited the final manuscript.

## Data availability statement

Data sharing is not applicable to this article as all created data is already contained within this article or in the supplementary material.

