## Supplementary Figure for "Sucrose transport and metabolism control carbon partitioning between stem and grain in rice"

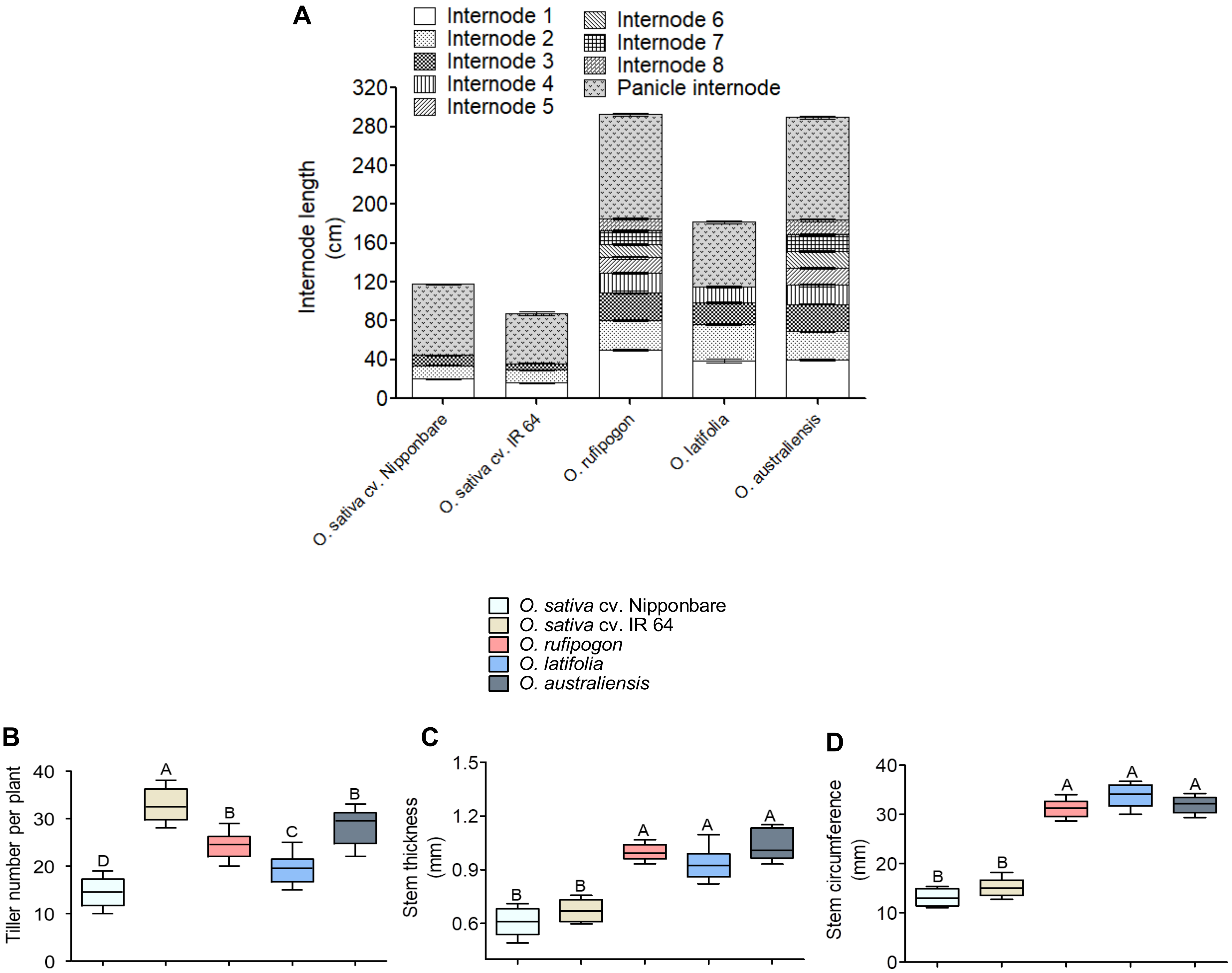


**Supplementary Fig. S1. Internode length, tiller number, and stem features of the selected cultivated and wild rice genotypes.**

Quantification of internode length (A), tiller number per plant (B), stem thickness (C), and stem circumference (D) of the selected rice genotypes. Each box and whisker plot shows the interquartile range with minimum and maximum values of ten data points from different plants. Significance of differences among the genotypes was calculated using One-Way ANOVA with Tukey's post-hoc test ($n = 10$, $p < 0.05$).

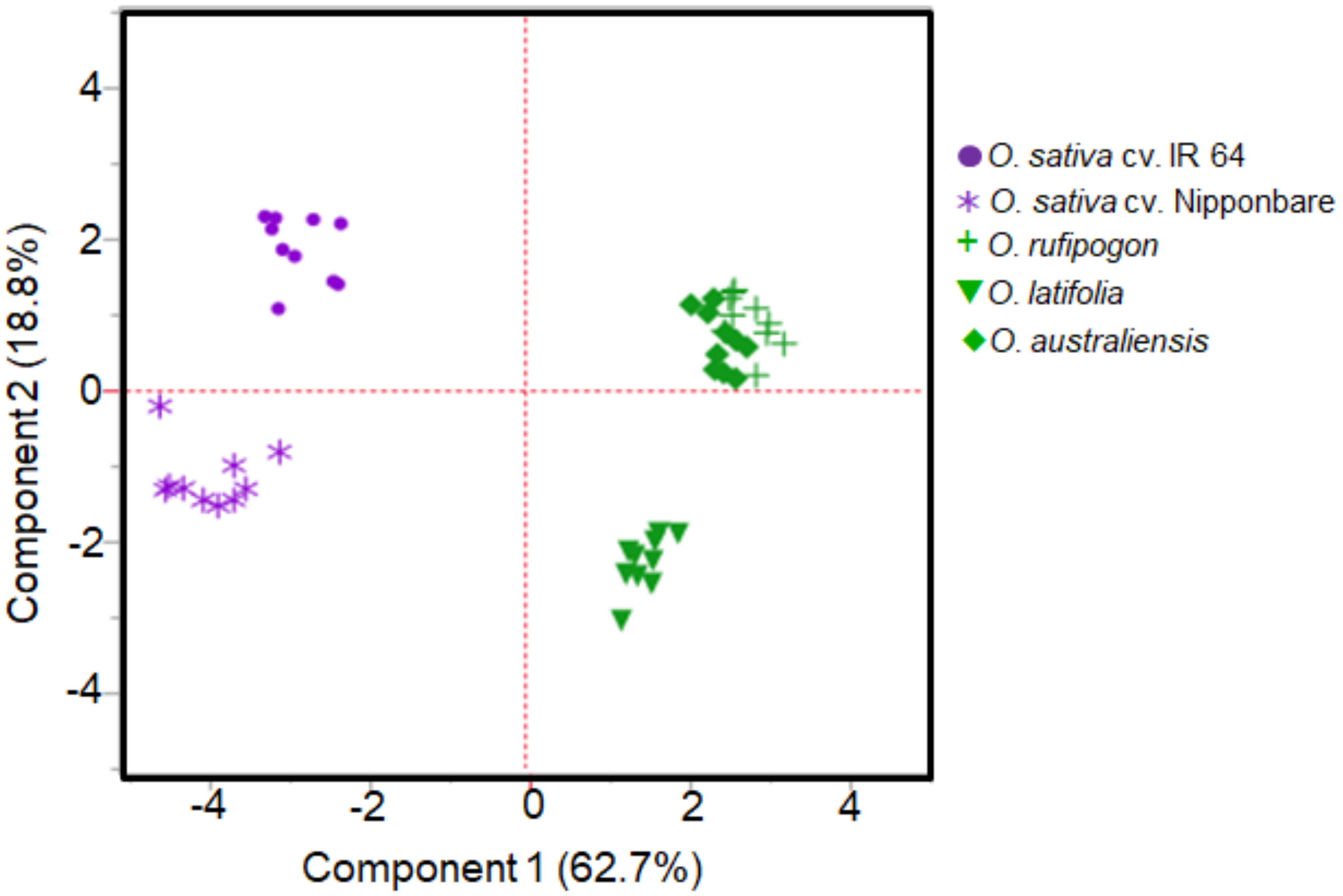


**Supplementary Fig. S2. Principal component analysis of biomass and yield traits of the selected cultivated and wild rice species.** Wild species are denoted in green color, and cultivated varieties in purple.

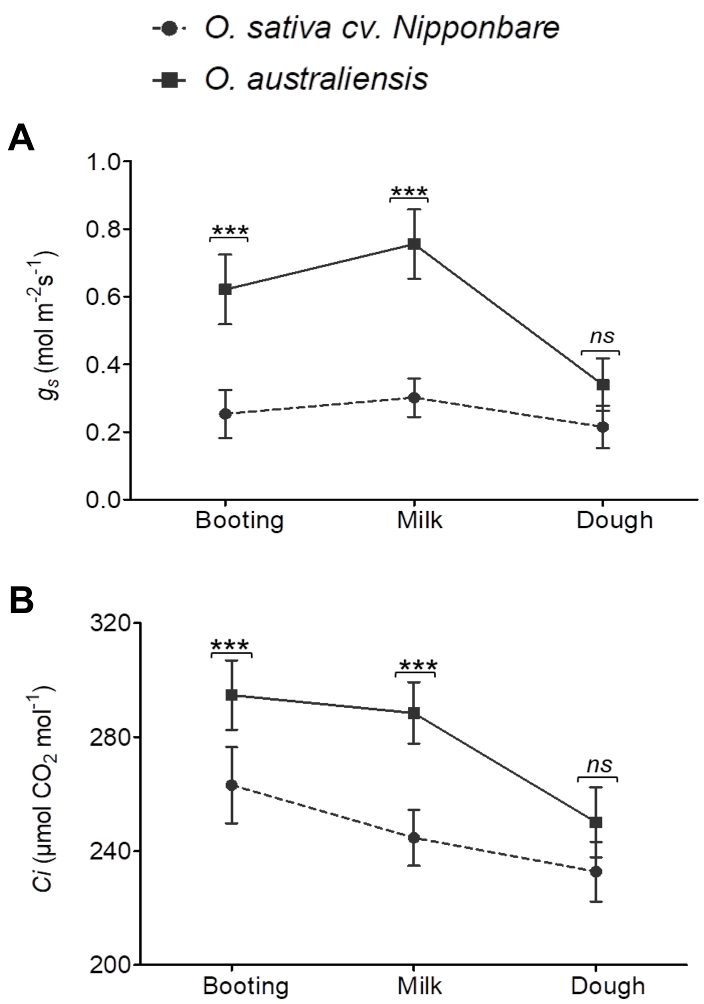


**Supplementary Fig. S3. Stomatal conductance ($g_s$) and intercellular $CO_2$ concentration (*Ci*) of a cultivated rice *O. sativa* cv. Nipponbare and a wild rice *O. australiensis* at different stages during booting and grain-filling.**
(A–B) Quantification of stomatal conductance ($g_s$, A) and intracellular $CO_2$ concentration (*Ci*, B) of flag leaves at booting-stage and milk- and dough- stage of grain filling of the selected cultivated and wild rice (n = 15). Data represents the mean and standard deviation (SD), and significance of differences between the genotypes was calculated using student t-test (*p < 0.05, **p < 0.01, ***p < 0.001).

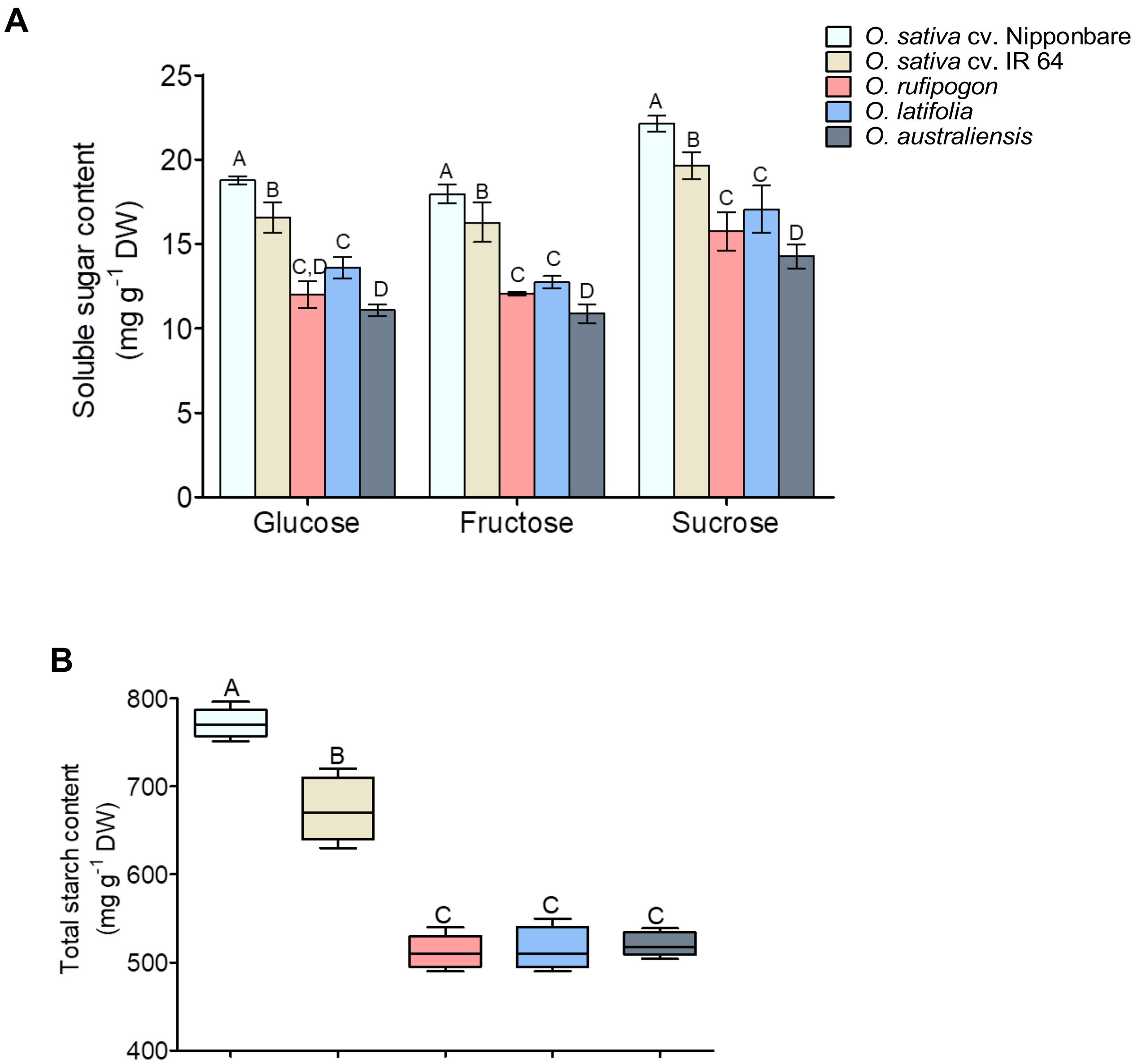


**Supplementary Fig. S4. Quantification of soluble sugars in matured seeds of the selected cultivated and wild rice genotypes.**

(A) Quantification of glucose, fructose, and sucrose content. Data represents the mean and standard deviation (SD) (n = 4). Significance of differences among the genotypes was calculated using One-Way ANOVA with Tukey's post-hoc test ($P \leq 0.05$). (B) Quantification of total starch content. Each box and whisker plot shows the interquartile range with minimum and maximum values. Significance of differences among the genotypes was calculated using One-Way ANOVA with Tukey's post-hoc test ($P < 0.05$).

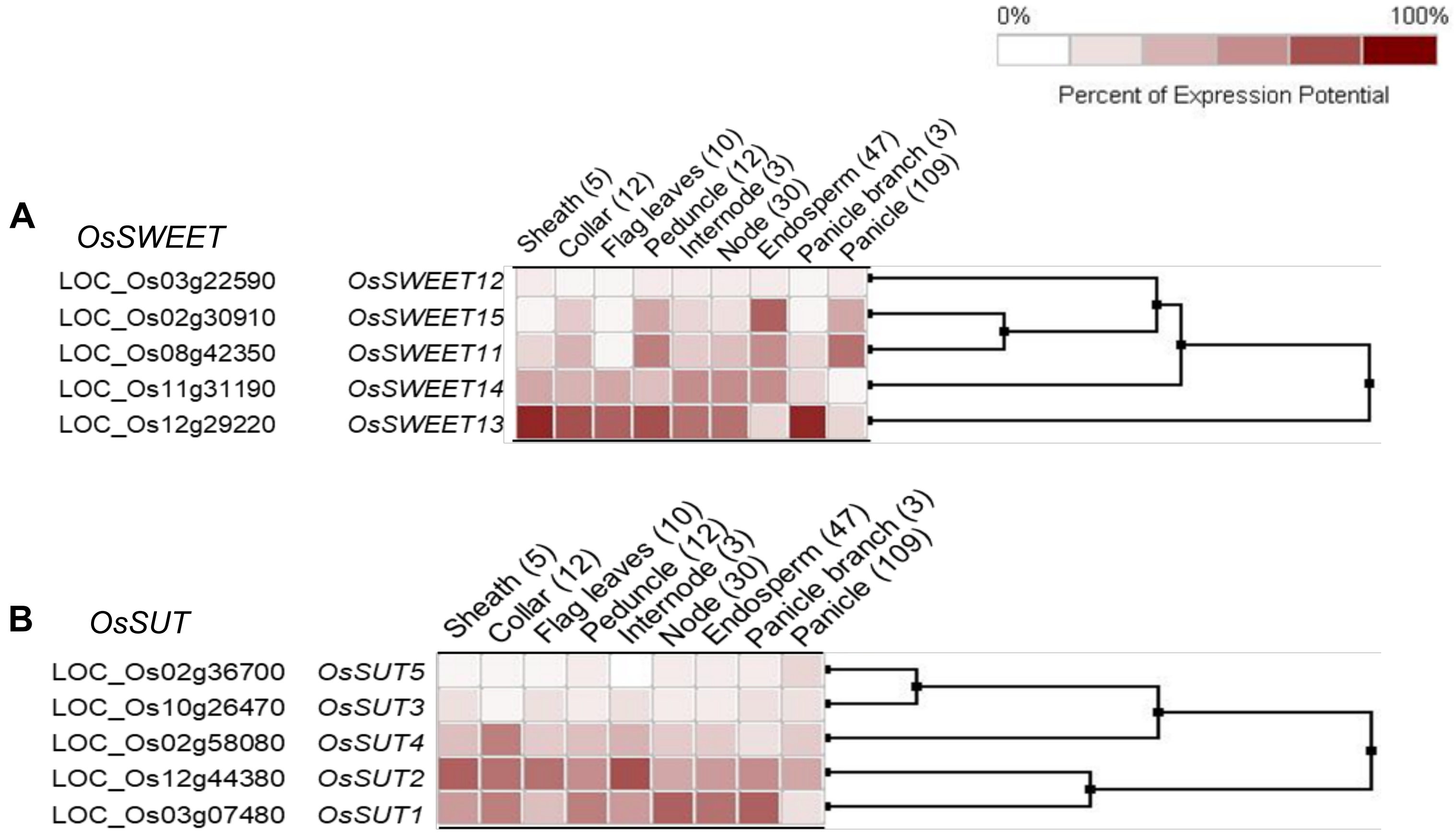


**Supplementary Fig. S5. *In silico* organ-specific expression analysis of rice genes encoding clade III SWEET (A) and SUT (B) transporters using publicly available data at Genevestigator (https://genevestigator.com/).**

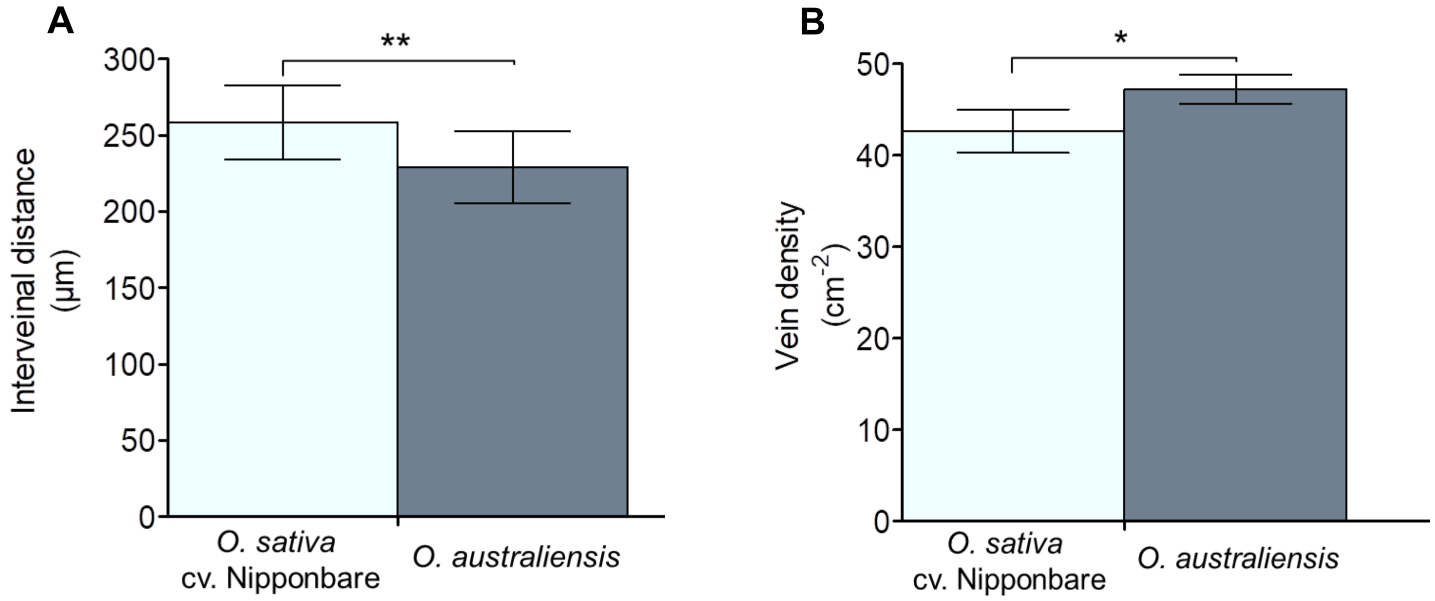


**Supplementary Fig. S6. Quantification of leaf interveinal distance (A) and vein density (B) for a cultivated rice *O. sativa* cv. Nipponbare and a wild rice *O. australiensis*.** Data represents the mean and standard deviation (SD) from fifteen cross-section images collected from five different plants. Significance of differences between the genotypes was calculated using student t-test (*p < 0.05, **p < 0.01, ***p < 0.001 ).

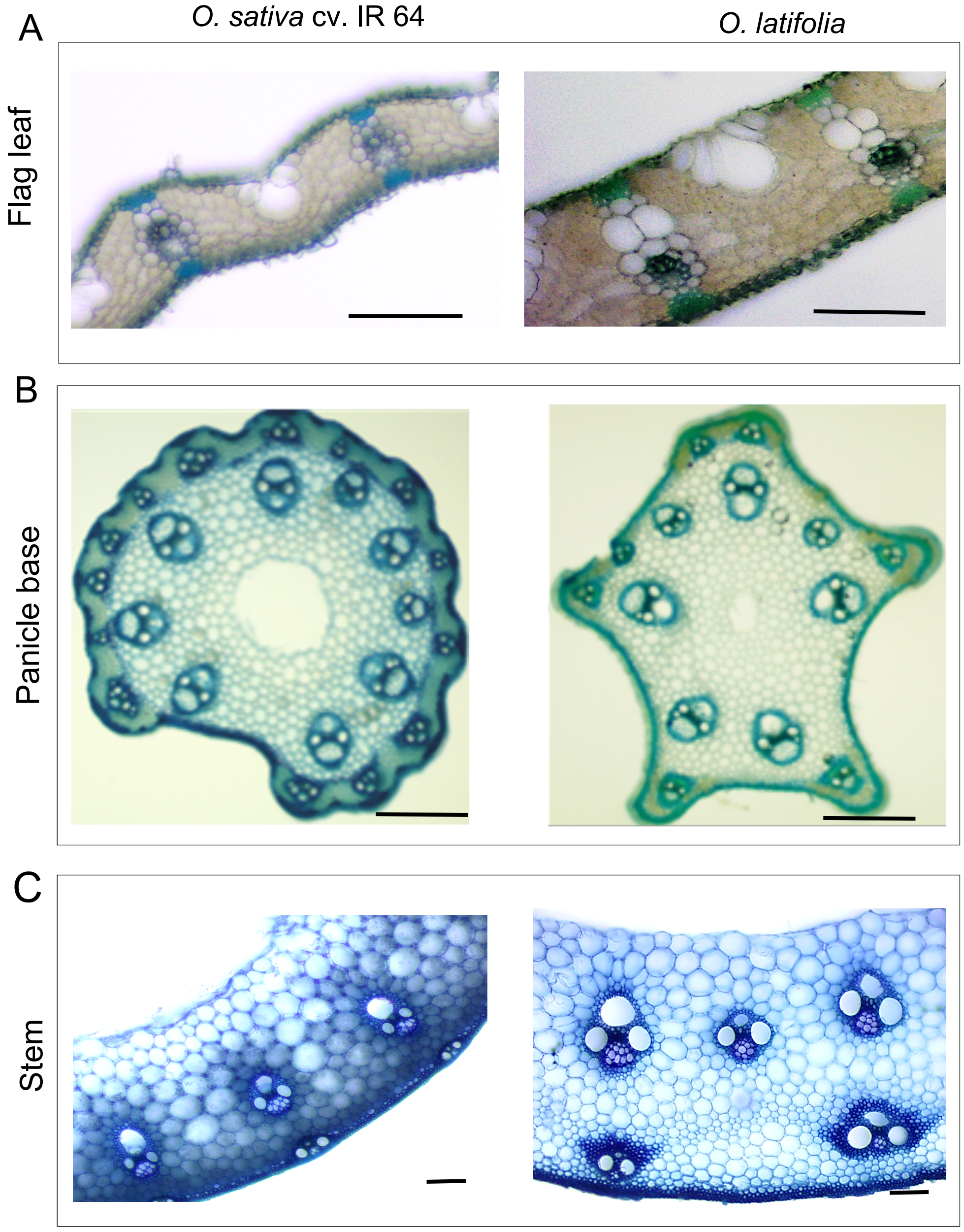


**Supplementary Fig. S7. Vascular features in flag leaf, panicle base, and stem of a cultivated rice *O. sativa* cv. IR 64 and a wild rice *O. latifolia*.**

(A) Cross-sections of flag leaves of the two species (scale bar = 100 μm).

(B) Transverse sections at the panicle base (0.5 cm above panicle node) of the two species (scale bar = 250 μm).

(C) Transverse sections of stems of the two species (scale bar = 100 μm).

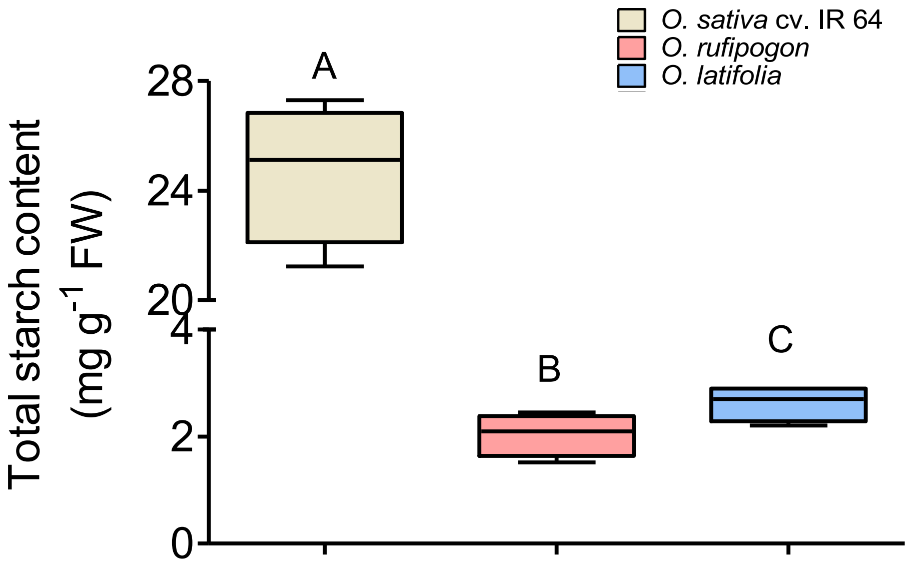


**Supplementary Fig. S8. Total starch content in the stem of a cultivated rice *O. sativa* cv. IR 64 and two wild rice species, *O. rufipogon* and *O. latifolia*, at milk-stage of grain-filling.** Each box and whisker plot shows the interquartile range with minimum and maximum values. Significance of differences among the genotypes was calculated using One-Way ANOVA with Tukey's post-hoc test ($n = 4$, $p \leq 0.05$).

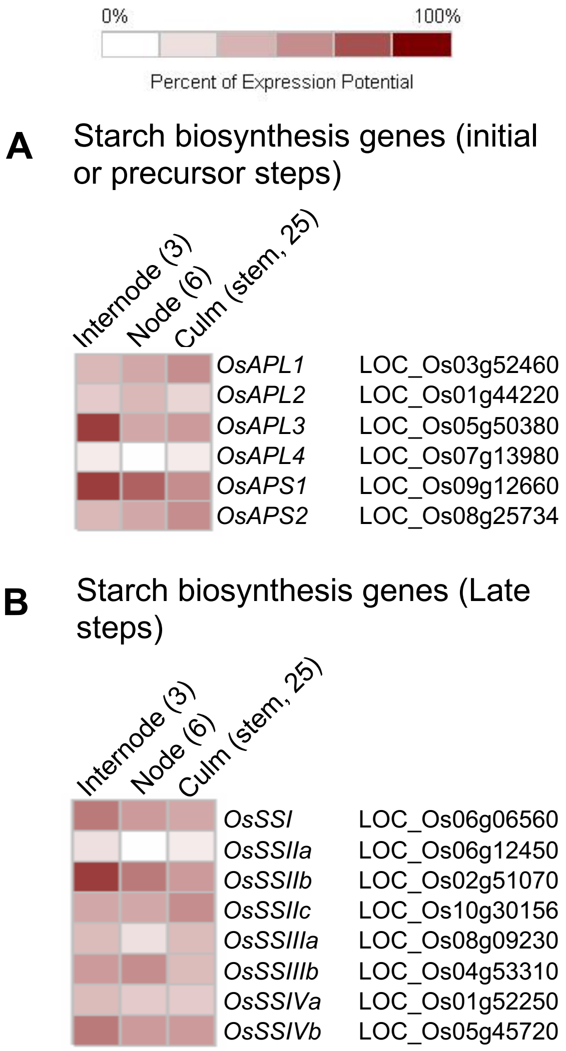


**Supplementary Fig. S9. *In silico* expression analysis of rice starch biosynthesis genes using publicly available data at Genevestigator (https://genevestigator.com/) in culm (stem), node, and internode.**

(A) Expression pattern of genes involved in initial or precursor steps of starch biosynthesis.

(B) Expression pattern of genes involved in late steps of starch biosynthesis.

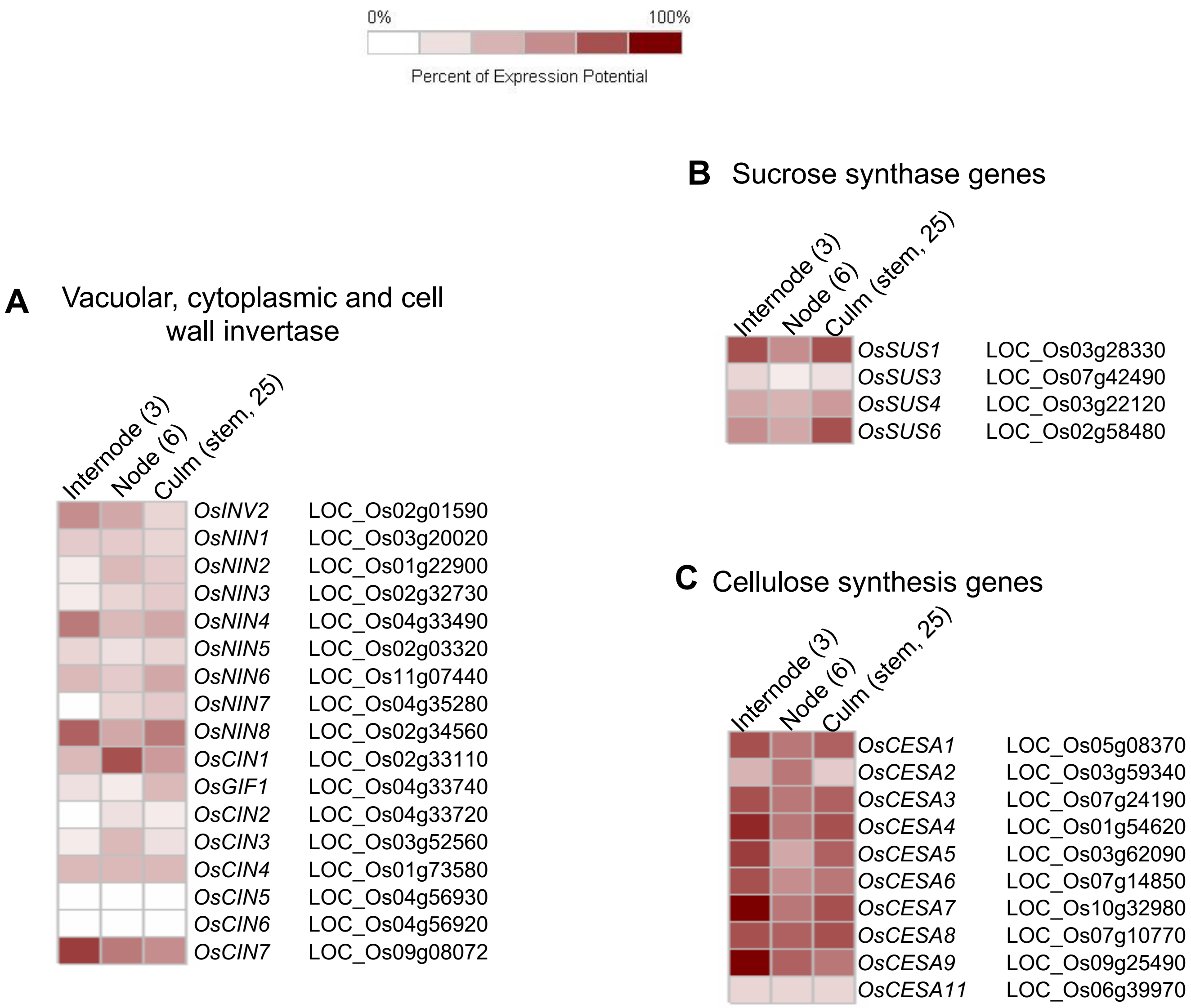


**Supplementary Fig. S10.** ***In silico*** **expression analysis of genes encoding rice invertases (A), sucrose synthases (B), and cellulose synthases (C) in culm (stem), node, and internode using publicly available data at Genevestigator (https://genevestigator.com/).**

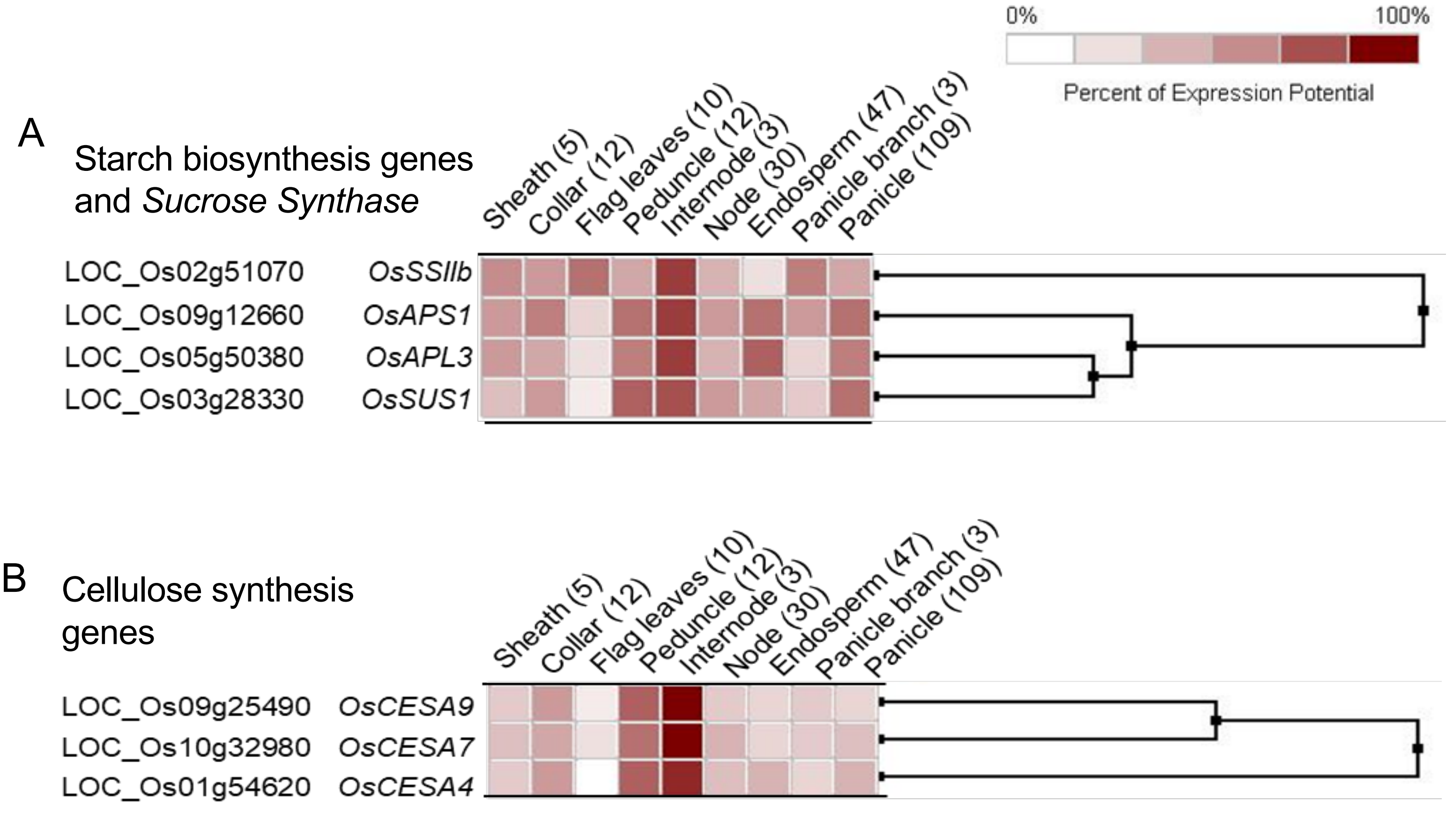


**Supplementary Fig. S11.** ***In silico*** **expression pattern of the selected key genes involved in sugar metabolism in rice using Genevestigator database (https://genevestigator.com/) across multiple different tissues.**
(A) Expression pattern of the selected starch-biosynthesis genes and *Sucrose Synthase1.*
(B) Expression pattern of the selected cellulose synthase genes

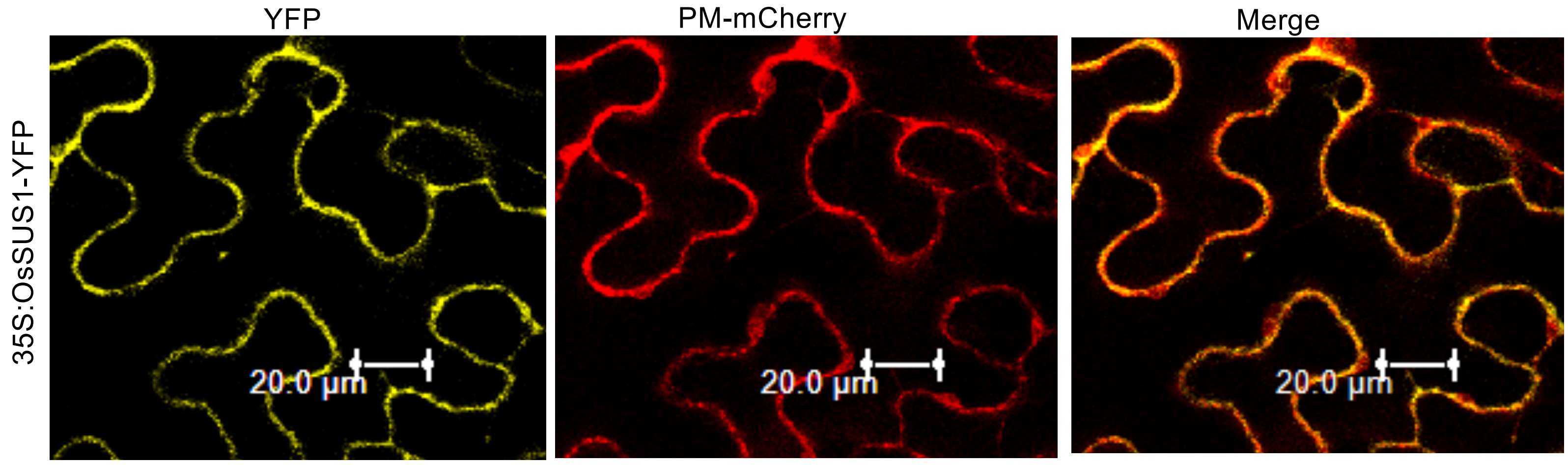


**Supplementary Fig. S12.** Localization of OsSUS1-YFP in the plasma membrane of leaf epidermal cells of *Nicotiana benthamiana*. Shown are the YFP signal (left column), plasma membrane marker PM-mCherry as RFP signal (middle column), and merged image of YFP and PM-mCherry in bright field background (right column).

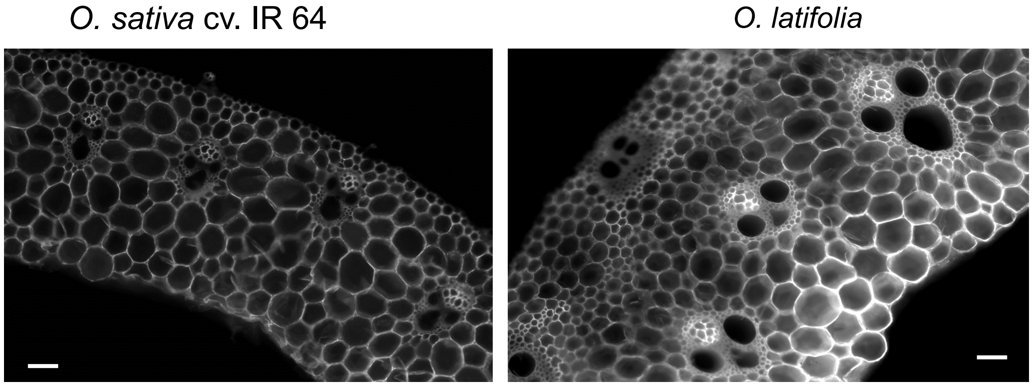


**Supplementary Fig. S13.** Calcofluor-white staining for cellulose deposition in the transverse stem sections of a cultivated rice *O. sativa* cv. IR 64 and a wild rice *O. latifolia*. Scale bar represents 500 µm.

**Supplementary Table S1.** List of primer pairs used in the study

| Gene | Forward (5'-3') | Reverse (5'-3') |
|---|---|---|
| **For qRT-PCR analysis** | | |
| *OsSWEET11* | TGGTTCTGCTACGGCCTCTT | GGTACCAGAAGTAGAGCCCCATCT |
| *OsSWEET12* | CGTGGAGTTCATGCCCTTCT | CGGCGTGGCAACGAA |
| *OsSWEET13* | CTACGCGCTGATCAAGTCCAA | GGGCGTAGGCGAGGTACAT |
| *OsSWEET14* | ATCTACTACGCGCTGCTCAAGTC | TAGACGAGGTAGACGGCGATGT |
| *OsSWEET15* | CCGTACGTGGTGACGCTCTT | ACGCACCCCACACCATTG |
| *OsSUT1* | TCATCCCTCAGGTGGTCATCG | CTTGGAGATCTTGGGCAGCAG |
| *OsSUT2* | GCATCAGCTGTGCCAACCT | CTGCTTCATCACTTCCAAAGGA |
| *OsSUT3* | TCCTCTTCGACACCGACTG | CAGCACGATCGAGTTAAGGAG |
| *OsSUT4* | ACAACGGTGTCCGAGAAGGT | GCACCCATCAGTCGGCATA |
| *OsSUT5* | TCGGCATGGTGTCCATGAG | CAATGGCAAGACCTTGGCC |
| *OsCIN1* | CGACCCTACCAA GTCTTCTCTTAG | CCCATTGTTGAAGACGTAAAGATG |
| *OsNIN8* | ATTGTGACAGGCTGTGATCC | TTCAACAGCCTTCTCTCAGC |
| *OsINV2* | CTTGGTACCTGTGCTAGATG | CTCGTAGATGGCTTTGGTG |
| *OsSUS1* | CATCTCAGGCTGAGACTCTGA | CAAATTCAATCGACCTTACTT |
| *OsAPL3* | TGGAAGGAACGTGGTCATCA | GTCGCGTTCTTCAGGATCAC |
| *OsAPS1* | CAGCACGAGTGTTCTTGGG | TAATTAGCACCCAACGGCAC |
| *OsSSIIb* | TGTTGCTGAACCGTTGGAAG | TTCATCACATTTGGCCCTGC |
| *CESA4* | GGTACCAACACCACAAGAAAG | GGACATAGCAGACATCTCTAC |
| *CESA7* | GAGGAGTTCCAGATCAAGAG | CTTCCGGTGAAGGAAGAGAG |
| *CESA9* | CTACATCAACACCACCATCTAC | GAGATGAAGAGCGCGATGAAG |
| *OsActin* | GAAGTGCGACGTGGATATTAG | CAGACACTGTACTTCCTTTCAG |
| *OsUbiquitin* | CTCGCCGACTACAACATCCA | TCTTGGGCTTGGTGTACGTCTT |
| **Localization study** | | |
| *OsSUS1* | ATGGGGGAAGCTGCCGGC | CTTGTTCGACGGCTCGCCC |
